# Identification of a secretion-enhancing *cis* regulatory targeting element (SECReTE) involved in mRNA localization and protein synthesis

**DOI:** 10.1101/303156

**Authors:** Osnat Cohen-Zontag, Lisha Qiu Jin Lim, Dvir Dahary, Tsviya Olender, Yitzhak Pilpel, Jeffrey E. Gerst

## Abstract

Earlier dogma states that mRNAs encoding secreted and membrane protein (mSMPs) reach the ER in a translation-dependent manner through the signal recognition particle (SRP) pathway. In this pathway, the signal sequence of the translation product is recognized by SRP and the mRNA-ribosome-nascent-chain-SRP complex is recruited to the ER via the interaction with an endoplasmic reticulum (ER)-localized SRP receptor. This model suggests that the translation product dictates the delivery of mRNAs to the ER and that the mRNA is a passive passenger. However, new evidence challenges this model and implies the existence of both translation - and SRP-independent mRNA localization to the ER, raising the possibility that mRNAs have an active role in determining their localization to the ER.

Besides serving as a template for protein translation, mRNAs carry information required for other regulatory processes such as mRNA processing, translation and transcription efficiency, degradation and localization. In yeast, mRNA localization governed by cis-acting sequence elements has been characterized for asymmetrically (*e.g.* bud) localized mRNAs that localize to, and are transported with, cortical ER. Now, we identify a *cis* motif in mSMPs that targets mRNAs mainly to the nuclear ER in yeast and increases both protein synthesis and secretion. Termed SECReTE, for secretion-enhancing *cis* regulatory targeting element, this motif was identified by computational analysis of genes encoding secretome proteins. SECReTE consists of ≥10 repetitive triplets enriched with pyrimidines (*i.e.* C’s and U’s) every third base (*i.e. NNY, N* - any nucleotide, *Y* - pyrimidine), and is found particularly in mRNAs coding for cell wall proteins. To study the physiological relevance of SECReTE, we introduced synonymous mutations that either elevate or decrease its overall score in genes coding for secreted proteins, without changing the protein sequence, and examined the physiological effects in yeast. An increase in the SECReTE score elevated the synthesis and secretion of endogenous proteins while, in contrast, a reduction led to less secretion and physiological defects. Importantly, the addition of SECReTE to the 3’UTR of an exogenous protein (*e.g.* SS-GFP) led to its increased secretion from yeast. SECReTE is present all through evolution and, thus, constitutes a novel RNA targeting motif found in both prokaryotes and eukaryotes.

## Introduction

mRNA targeting and localized translation is an important mechanism that provides spatial and temporal control of protein synthesis. The delivery of mRNA to specific subcellular compartments has a major role in the establishment of polarity in various of organisms and cell types, and was shown to be crucial for the proper function of the cell^1,2^ Interestingly, the localization of mRNA is often governed by *cis*-acting elements (zipcodes) embedded within the mRNA sequence^1,2^, RNA binding proteins (RBPs) recognize this sequence and act together with molecular motors to direct the mRNA to its final destination.

The endoplasmic reticulum (ER) is the site of synthesis of secreted and membrane (SMP; secretome) proteins. According to dogma, mRNAs encoding for SMPs (mSMPs) are delivered to the ER by a distinct translation-dependent mechanism, termed the signal recognition particle (SRP) pathway^3-5^. According to this model, protein translation begins in the cytoplasm and when SMPs undergo translation, a signal peptide present at their amino terminus emerges from the exit channel of translating ribosome and is recognized by the SRP. The SRP then is recruited to its receptor on the ER membrane and translocation of ribosome-mRNA-nascent polypeptide chain complex from the cytoplasm to the ER occurs. There, translating ribosomes interact with the translocon to enable co-translational protein translocation and mRNA anchoring^6,7^. Thus, the SRP model describes mSMPs as components with no active role in the ER translocation process.

However, multiple lines of evidence suggest that there are additional pathways for the delivery of mRNAs to the ER^8,9^. First, loss of the SRP pathway did not result in lethality of yeast^10^ and mammalian cells^11^, and also did not have a significant effect upon membrane protein synthesis and global mRNA distribution between the cytoplasm and the ER^11^. Second, genome-wide analyses of the distribution of mRNAs encoding soluble and membrane proteins between cytosolic polysomes and ER-bound polysomes have demonstrated a significant overlap in the composition of the mRNA in the two fractions and showed that cytosolic protein-encoding mRNAs are broadly represented on the ER^12-16^. This means that mRNAs lacking an encoded signal sequence can localize to the ER. In agreement with these findings, removal of the signal sequence and the inhibition of translation did not disrupt mSMP localization to the ER^14,17,18^. Third, subsets of secretome proteins are known to localize to the ER in an SRP-independent pathway^19,20^. These proteins are thought to translocate into the ER after translation in the cytosol^21^. In a study that utilized a technique for a specific pull-down of ER-bound ribosomes^22^, it was found that there is no significant difference in the enrichment of mRNAs encoding SRP-dependent proteins in comparison to mRNAs encoding SRP-independent proteins on ER membranes. In addition, a subset of ribosomes managed to reach the ER before the emergence of the signal sequence. A possible explanation for these observations could be that mRNAs reach the ER before the ribosomes in an SRP-independent mechanism. If mRNA targeting to the ER does not begin until signal peptide emergence, membrane-bound ribosomes should not be translating portions of transcript upstream of the signal peptide. However, this was not the case, as translating membrane-bound ribosomes were found evenly distributed across entire transcripts^23^. This suggests that these mRNAs may localize to ER prior to translation initiation.

Although it has been difficult to identify clear *cis*-elements within mRNA that direct it to the Er^24-27^, specific sequence characteristics of mSMPs have been identified. For example, sequence analysis of the region encoding the signal sequence revealed a low usage of adenine to create no-A stretches within this sequence^28^. Additionally, mRNAs encoding membrane proteins have a high degree of uracil enrichment, as well as pyrimidine usage, in comparison to mRNAs encoding cytosolic proteins^8,26,29,30^. These findings raise the possibility that the motif resides in a general, more diffuse, fashion in the sequence composition of the mRNA molecule.

By examining the sequences of transmembrane domain (TMD)-containing regions in mRNAs, we have now identified high content stretches of pyrimidine (C and U) repeats every third base (*NNY, N* - any nucleotide, *Y* - pyrimidine) that can be ≥10 nucleotide triplets in length. Analysis of the transcriptomes of several eukaryotic organisms (*e.g.*, human, mouse, fish, fly, and yeast), revealed that this sequence pattern is significantly over-represented in mRNAs encoding for secretome proteins, which presumably localize to the ER. The location of the motif is not restricted to the coding region but can be present in the untranslated regions (UTRs). Although we originally found the motif by analyzing the sequences of TMDs in secreted membrane proteins, in fact it is enriched at a higher level in mRNAs encoding secreted proteins that lack TMDs. Both computational and experimental tools were utilized to establish the existence and significance of this motif. Computational analysis verified that mSMPs are the group most enriched with the motif, while synonymous mutations that either elevated or decreased motif strength (*i.e.* number of consecutive pyrimidine repeats) in mRNAs encoding yeast invertase, *SUC2*, as well as cell wall proteins, *CCW12* and *HSP150*, enhanced or reduced protein synthesis and secretion, respectively. This motif, which could serve as a signal for mRNA localization and translation at the ER, we have named the <u>s</u>ecretion-<u>e</u>nhancing *<u>c</u>is* <u>re</u>gulatory <u>t</u>argeting <u>e</u>lement (SECReTE). SECReTE is found in prokaryotes and in both lower and higher eukaryotes. This suggests that it may have a conserved role in the translational control of mRNAs either as a targeting motif or in other processes such as translation efficiency, mRNA processing (*i.e.*, polyadenylation, capping, splicing), mRNA decay, and secondary structure, *etc.* We propose that SECReTE is important, not only to understand how mRNAs may reach the ER, but may have practical applications in the field of biotechnology.

## Materials and Methods

### Yeast strains and plasmids used

The yeast strains and plasmids used in this study are listed in Table S1 and S2, respectively.

### Yeast strains, genomic manipulations, and growth conditions

Yeasts were grown at the indicated temperature either in a standard growth medium (1% Yeast Extract, 2% Peptone, 2% Dextrose) or synthetic medium containing 2% glucose [*e.g.*, synthetic complete (SC) and selective SC dropout medium lacking an amino acid or nucleotide base]^31^. Deletion strains using the *NAT* antibiotic resistance gene in WT (BY4741) cells were created using standard LiOAc transformation procedures and with nourseothricin (100μg/ml) for selection on synthetic solid medium. For the creation of SECReTE mutant strains, SECReTE gene fragments were designed with the appropriate modifications, from the first to the last mutated base, and synthesized either as a gBlock^®^ (Integrated DNA Technologies, Inc., Coralville, IA, USA) or cloned into a pUC57-AMP vector (Bio Basic Inc.). Both (-)SECReTE and (+)SECReTE strains were generated. *SUC2*(-)SECReTE, *SUC2* (+)SECReTE and *CCW12*(-)SECReTE strains were constructed in the BY4741 background genome using the *delitto perfetto* method for genomic oligonucleotide recombination^32^, in which the CORE cassette from pGKSU^32^ was integrated first into the genomic region corresponding to site of the SECReTE gene fragment. The CORE cassette contains the *URA3* selection marker with an *I-SceI* homing endonuclease site and a separate inducible *I-SceI* gene. The SECReTE gene fragment for *CCW12*(-)SECReTE was amplified from the synthetic gBlock using primer sequences containing 20 bases of homology to both the region outside of the desired genomic locus and the CORE cassette. The amplified SECReTE gene fragment subsequently replaced the CORE cassette in the desired genomic site through an additional step of integration. CRISPR/Cas9 was utilized instead to generate the *HSP150* mutant strains. *HSP150*(-)SECReTE and *HSP150*(+)SECReTE were created in the BY4741 genome. The CRISPR/Cas9 procedure involved deletion of the native genomic region corresponding to the SECReTE gene fragment, using the *NAT* cassette from pFA6-NatMX6. A CRISPR/Cas9 plasmid vector was designed to express the Cas9 gene, a guide RNA that targets the *NAT* cassette, and the *LEU2* selection marker. The CRISPR/Cas9 plasmid was co-transformed with the amplified SECReTE gene fragment to replace the *NAT* cassette. Standard LiOAc-based protocols were employed for transformations of plasmids and PCR products into yeast. Transformed cells were then grown for 2-4 days on selective media. Correct integrations were verified at each step using PCR and, at the final step, accurate integration of the (-)SECReTE or (+)SECReTE sequences was confirmed by DNA sequencing.

### Quantitative RT-PCR (qRT-PCR)

RNA was extracted and purified from overnight cultures using a MasterPure Yeast RNA Purification kit (Epicentre Biotechnologies). For each sample, 2μg of purified RNA was treated with DNase (Promega, Madison, WI, USA) for 2hrs at 37°C and subjected to reverse transcription (RT) using Moloney murine leukemia virus RT RNase H(-) (Promega) under the recommended manufacturer conditions. Primer pairs were designed, using NCBI Primer-Blast^33^, to produce only one amplicon (60-70bp). Standard curves were generated for each pair of primers and primer efficiency was measured. All sets of reactions were conducted in triplicate and each included a negative control (H_2_O). qRT-PCR was performed using a LightCycler^®^480 device and SYBR^®^ Green PCR Master Mix (Applied Biosystems^®^, Waltham, Massachusetts, USA). Two-step qRT-PCR thermocycling parameters were used as specified by the manufacturer. Analysis of the melting curve assessed the specificity of individual real-time PCR products and revealed a single peak for each real-time PCR product. The *ACT1* or *UBC6* RNAs were used for normalization and fold-change was calculated relative to WT cells.

### Drop test growth assays

Drop test assays were performed by growing yeast strains in YPD medium to mid-log phase and then performing serial dilution five times (10-fold each) in fresh medium. Cells were spotted onto plates with different conditions and incubated for 48hrs, prior to photo-documentation. Calcofluor White (CFW) or Hygromycin B (HB) sensitivity was tested by spotting cells onto YPD plates containing either 25μg/ml HB or 50μg/ml CFW (dissolved in DMSO, prepared as described^34^), following the protocol as mentioned above.

### Hsp150 and GFP secretion assays

For the induction of Hsp150 secretion, strains were grown in YPD overnight at 26°C, diluted in YPD medium to 0.2 O.D._600_ units, and then incubated at 37°C and grown until log-phase. For GFP secretion, yeast were grown O/N to 0.2 O.D._600_ at 30°C in synthetic selective medium containing raffinose as a carbon source, then diluted to 0.2 OD O.D._600_ units in YP-Gal and grown to mid-log phase (0.6-0.8 O.D._600_) at 30°C. Next, 1.8ml of the culture was taken from each strain and centrifuged for 3mins at 1900 x *g* at room temperature. Trichloroacetic Acid (100% w/v) protein precipitation was performed on the supernatant and protein extraction, using NaOH 0.1M, was performed on the pellet ^35^. Samples were separated on SDS-PAGE gels, blotted electrophoretically onto nitrocellulose membranes, and detected by incubation with rabbit anti-Hsp150 [1:10,000 dilution; gift from Jussi Jäntti (VTT Research, Helsinki)] or monoclonal mouse anti-GFP (Roche Applied Science, Penzberg, Germany) antibodies followed by visualisation using the Enhanced Chemiluminescence (ECL) detection system with anti-rabbit peroxidase-conjugated antibodies (1:10,000, Amersham Biosciences). Protein markers (ExcelBand 3-color Broad Range Protein Marker PM2700, SMOBiO Technology, Inc., Hsinchu, Taiwan) were used to assess protein molecular mass.

### Invertase assay

Invertase secretion was measured as described previously^36^. Cell preparation for the invertase assay was performed as described in ^37^. The protocol was optimized based on previous work^38^. Internal and external activities were expressed in units based on absorption at 540 nm (1 U = 1 μmol glucose released/min per OD unit).

### Single-molecule FISH

Yeast cells expressing Sec63-GFP were grown to mid log phase and shifted to low glucose-containing medium [0.1% glucose] for 1.5 h to induce *SUC2* expression. Cells were fixed in the same medium upon the addition of formaldehyde (3.7% final concentration) for 45min. Cells were gently washed twice with 0.1M potassium phosphate buffer, pH 7.4 containing 1.2M sorbitol, after which cells were spheroplasted in 1ml of freshly prepared spheroplast buffer [0.1M potassium phosphate buffer, pH 7.4, 1.2M sorbitol, 20mM ribonucleoside vanadyl complexes (Sigma-Aldrich, St. Louis, MO), 1×Complete Protease Inhibitor Cocktail, 28mM β-mercaptoethanol, 120U/ml RNasin Ribonuclease Inhibitor, and Zymolase (10 kU/ml)] for 30min at 30°C. The spheroplasts were centrifuged for 4min at 1300 × *g* at 4°C and washed twice in 0.1M potassium phosphate buffer, pH 7.4 containing 1.2M sorbitol. Spheroplasts were then resuspended in 70% ethanol and incubated for 1hr at 4°C. Afterwards, cells were centrifuged at 1300 × *g* at 4°C for 4min, washed with WASH buffer (0.3M sodium chloride, 30mM sodium citrate, and 10% formamide), and incubated overnight at 30°C in the dark with a hybridization mixture containing 0.3M sodium chloride, 30mM sodium citrate, 10% dextran sulfate, 10% formamide, 2mM ribonucleoside vanadyl complexes, and the TAMRA-labeled Stellaris probe mix for *SUC2* (Biosearch Technologies, Novato, CA). After probe hybridization, labeled spheroplasts were centrifuged at 1300 × g, the hybridization solution aspirated, and the spheroplasts incubated for 30min at 30°C in WASH buffer. Cells were then centrifuged and resuspended in a solution containing 0.3M sodium chloride and 30mM sodium citrate. *SUC2* mRNA co-localization with the ER was visualized using a DeltaVision imaging system (Applied Precision, Issaquah, WA, USA). Images were processed by deconvolution.

### Computational Analyses

#### Gene ontology

Definition of the secretome was used according to Ast *et al.*^19^. This group includes all genes that contain at least one TMD and/or signal sequence and are not mitochondrial. TMHMM tool was used to define TMD-containing proteins. Cell wall and tail anchored genes were defined according to UniProt. Data from Jan *et al*^22^ was used to define other groups of genes and for defining human GO terms. The GO Slim Mapper tool (SGD) (www.yeastgenome.org/cgi-bin/GO/goSlimMapper.pl) was used to classify SECReTE10-and SECReTE15-positive genes.

#### Permutation test analysis

For permutation analysis, each gene sequence was randomly shuffled 1000 times and the SECReTE was scored for each of the shuffled sequences. To evaluate the significance of the ability of SECReTE to appear randomly, a Z score was calculated for each gene according to the formula: Z= (Observed - mean)/STD. Observed is the value that was measured from the gene sequence. (*e.g.* SECReTE score for the gene). Mean is the average SECReTE score for all shuffled sequences of the gene. STD is the standard deviation of the SECReTE score from all shuffled sequences of the gene.

#### Identification of cell wall motif

Motif search was performed by MEME suites^39^, at http://meme-suite.org/tools/meme.

## Results

### Identification of a pyrimidine repeat motif in mRNAs encoding yeast secretome proteins

Because codons encoding hydrophobic residues are enriched in pyrimidines in their second position^30^, we examined mRNAs encoding secretome proteins in yeast for the presence of consecutive pyrimidine repeats every third nucleotide (*i.e. YNN, NYN*, or *NNY*) in the coding and UTR regions. First, we determined how many such repeats might best differentiate secretome protein-encoding mRNAs from non-secretome protein-encoding mRNAs. For that, the number of repeats along an mRNA transcript was scored according to a defined threshold (*e.g.* 5, 7, 10, 12, and 15 repeats). For a random motif we expected to see a linear correlation between the probability of its appearance(s) in a gene and gene length, as shown in Figure 1A. We operationally employed SECReTE scores between 5 and 15 (*e.g.* 5,7,10,12,15) and observed a direct correlation between SECReTE number and gene length for SECReTE5 and SECReTE7 (Figure 1D). However, the dependency on gene length becomes significantly weakened above SECReTE10 (Figure 1A). This implies the presence of ≥10 consecutive repeats is not a random phenomenon and may be important.

**Figure 1.**
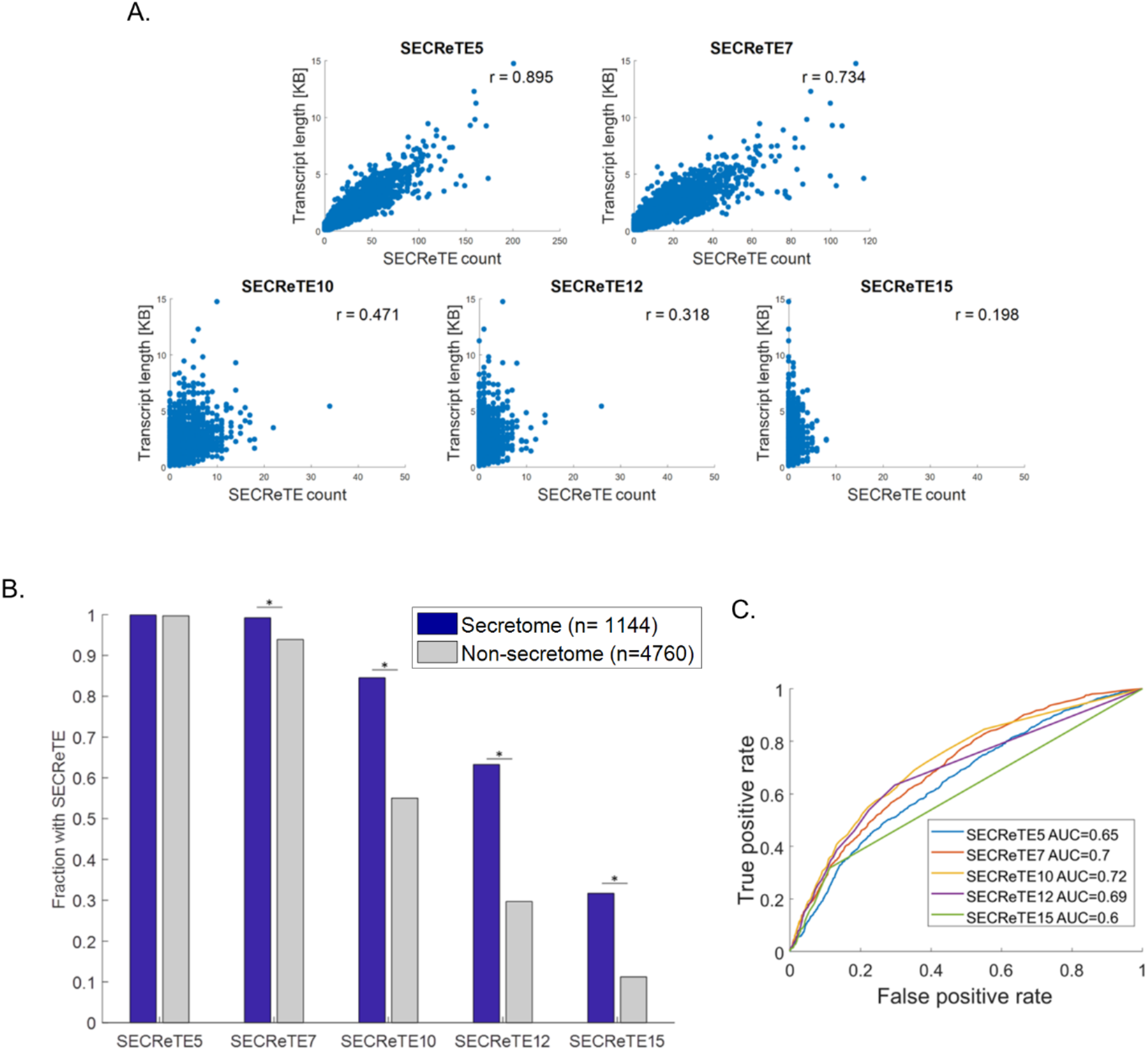
Determination of the number of *NNY* repeats to use as a threshold for SECReTE. **(A) Correlation between SECReTE number and transcript length.** The total SECReTE score was calculated for each yeast gene (5904 scored) by counting the number of consecutive *NNY* repeats present in the transcript sequence according to the indicated threshold, and in all three frames. Scatter plots represent the correlation between the SECReTE score and gene length. The SECReTE score does not correlate with gene lengths above a threshold of 10 NNY repeats (SECReTE10). R score represents the Pearson correlation coefficient. **(B) SECReTE motifs are more abundant in the mRNAs coding for secretome proteins than for non-secretome proteins.** SECReTE presence, according to the indicated threshold, was scored in mRNAs coding for secretome (blue) and nonsecreted (gray) proteins. Bars represent the fraction of SECReTE positive transcripts at the indicated threshold. SECReTE abundance is significantly higher in secretome mRNAs. \**p*≤ 2.28E-13. **(C) SECReTE10 maximizes the ability to distinguish secretome transcripts.** ROC curves were plotted for each of the indicated thresholds. Secretome transcripts were used as the “true positive” set, while non-secretome transcripts were used as the “true negative” set. The AUC (area under the curve) of SECReTE10 was the highest.

If SECReTE repeats above 10 (*e.g.* SECReTE10) play a role in protein secretion, we expect them to be more abundant in mRNAs encoding secretome proteins, as defined according to Ast *et al.*^19^. To test this possibility, we divided the complete yeast genome into two groups: secretome and non-secretome, and calculated the fraction of transcripts that contain SECReTE in each group. We found transcripts coding for secretome proteins are enriched with SECReTE motifs >7 (Figure 1B), as opposed to transcripts encoding non-secretome proteins. To test the number of repeats that give the most significant separation between secretome and non-secretome transcripts, we evaluated the different thresholds for their ability to classify mSMPs using receiver operator characteristics (ROC) analysis. *Bona fide* secretome protein-encoding transcripts were used as a true positive set and non-secretome protein-encoding transcripts were defined as true negatives. As seen (Figure 1C), the SECReTE10 threshold maximally differentiated secretome transcripts from non-secretome transcripts. As SECReTE10 did not show a dependency upon gene length and gave the most significant separation between secretome and non-secretome transcripts, we used it as the threshold by which to define motif presence in subsequent analyses. Previous studies have used high throughput analyses to quantify the level of enrichment of transcripts on yeast ER-bound ribosomes and ER membranes^22,23^. By comparing the cumulative distribution of the ER enrichment value of SECReTE10-containing transcripts to transcripts lacking SECReTE10, we could verify that a higher fraction of SECReTE10-containing transcripts is indeed enriched on ER-bound ribosomes (Figures S1A-B) and ER membranes (Figure S1C). In contrast, SECReTE10-containing transcripts are not enriched on mitochondrial ribosomes, in comparison to transcripts lacking SECReTE10 (Figure S1D).

### SECReTE abundance in mSMPs is not dependent on the presence of a TMD

TMDs encoding mRNA sequences are enriched with uracil (U), mainly in the second position of the codon (*NYN*)^29,30^. Since most secretome proteins contain TMDs, their presence alone might be the reason for motif enrichment in secretome transcripts. To ascertain whether SECReTE enrichment in mSMPs is not merely due to the presence of encoded TMDs, we determined at which position of the triplet the pyrimidine (*Y*) is located in the SECReTE10 elements: first (*YNN*); second (*NYN*); or third (*NNY*). We calculated SECReTE10 abundance separately for each position using only the coding sequences (*i.e.* from start codon to the stop codon) and without the UTRs. While the signal is present in the second position (Figure 2A; *NYN*), as expected, it is also abundant in the third position of the codon (Figure 2A; *NNY*). The latter finding implies that the TMD may not be the only factor that affects SECReTE enrichment in mSMPs. In contrast, the SECReTE10 element is poorly represented in the first position (Figure 2A; *YNN*).

**Figure 2.**
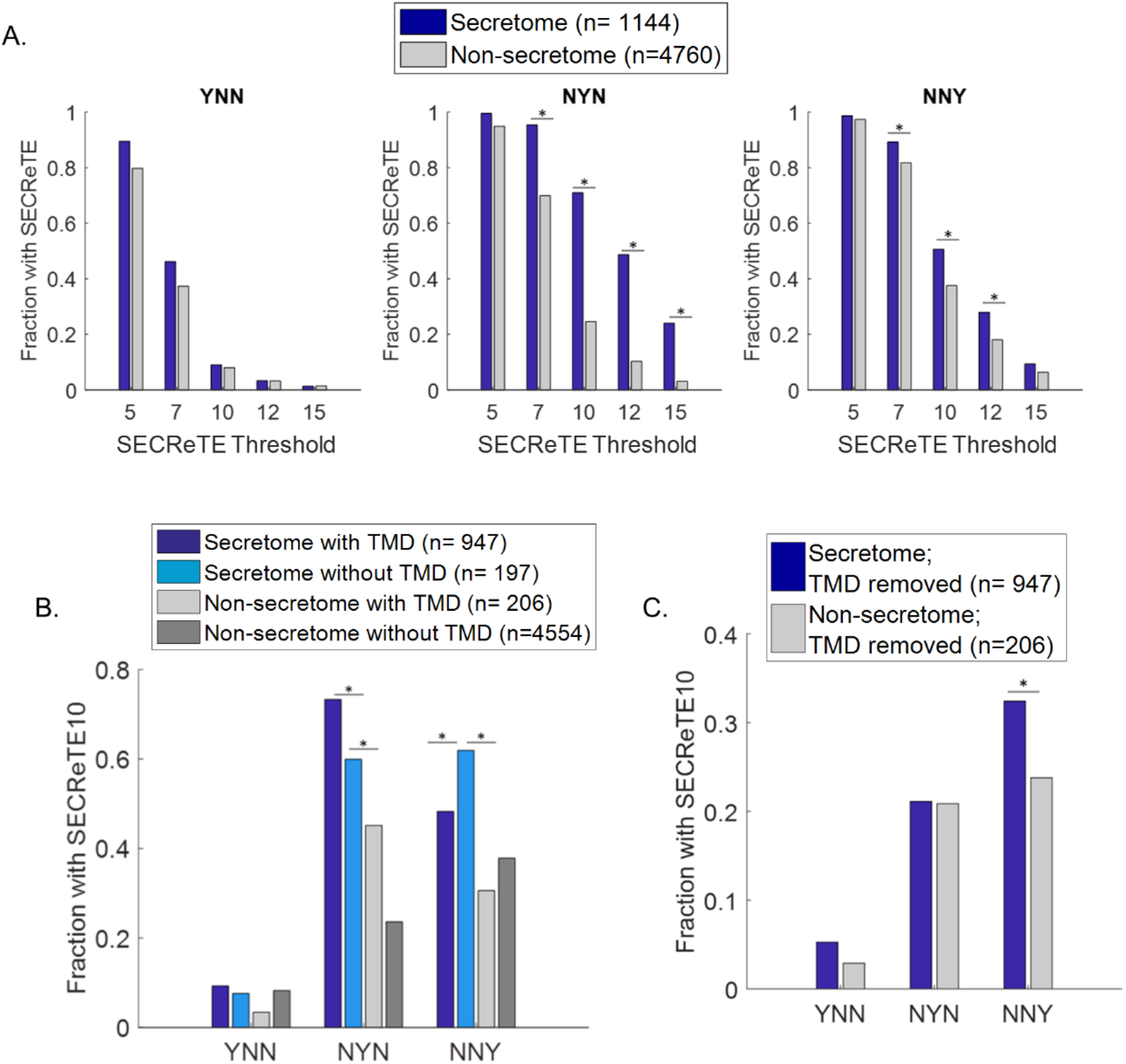
SECReTE abundance in mSMPs is TMD-independent. **(A) SECReTE is abundant in the second position of the codon.** SECReTE abundance was calculated for each codon position separately. SECReTE abundance in mSMPs is most significant in the second codon position, but significant differences were also detected in in the third position. \**p*≤ 9.9E-10. **(B) SECReTE is also highly abundant in the mRNAs encoding soluble secretome proteins.** SECReTE10 presence was examined separately for TMD-containing proteins and soluble secreted proteins. A higher fraction of mRNAs coding for soluble secreted proteins (Secretome without TMD; cyan) contains SECReTE in comparison to non-secretome transcripts, either with or without a TMD (Non-secretome with TMD; dark gray, Non-secretome without TMD; light gray). In the third codon position (*NNY*), the fraction of soluble secreted proteins is even larger than TMD-containing secretome proteins and is significant. \**p*≤3.03E-3. **(C) SECReTE is abundant at the third position after removal of the TMD sequence.** SECReTE10 presence was scored in mRNAs coding for membrane proteins after the encoded TMD was removed. SECReTE10 is significantly more abundant in the third position (*NNY*) in mRNAs encoding secretome proteins (blue) than non-secretome proteins (gray), even after removal of the TMD sequence. \**p* =0.01.

Next, we checked for the presence of SECReTE10 in mRNAs coding for TMD-containing proteins and soluble secreted proteins separately. As expected, more transcripts encoding TMD-containing secretome proteins contain SECReTE10 in the second position (*NYN*) than transcripts coding for soluble secreted proteins (Figure 2B). However, the fraction of SECReTE10-containing transcripts coding for soluble secreted proteins in the third position (*NNY*) is even higher. This provides compelling evidence for SECReTE10 enrichment in transcripts independent of the encoded TMD regions. Correspondingly, when the TMD was artificially removed from the sequences of mRNAs encoding membrane proteins, the secretome genes were no longer enriched with second position SECReTE10s (Figure 2C; *NYN*), although, the enrichment of SECReTE10 at the third position remained highly abundant (Figure 2C; *NNY*).

### SECReTE abundance is not dependent upon codon composition

There is a possibility that SECReTE enrichment results from codon composition of the transcript. To check this possibility, we performed permutation test analysis. In this case, each gene sequence was randomly shuffled x 1000, while codon composition remained constant. We then calculated the Z-score (*i.e.* number of standard deviations from the mean) of SECReTE10 for each gene to evaluate the probability of the signal to appear randomly. By looking at Z-score distribution in secretome and non-secretome genes, it can be concluded that SECReTE enrichment in mSMPs is not a random phenomenon and is not dependent on codon composition (Figure S2A). This conclusion is valid for mSMPs encoding both membranal and soluble proteins (Figure S2B). We also conducted the analysis for each codon position separately. For that, we calculated the fraction of genes with a significant Z-score (≥1.96) for each position separately. The fraction of genes with a significant Z-score was larger in secretome genes than in the non-secretome genes at both the second and third positions of the codon (Figure S2C), strengthening the notion that SECReTE is significantly more enriched in those positions. This finding is not dependent on the presence of TMDs, since the fraction of genes with a significant Z-score was larger for both soluble and TMD-containing secretome transcripts, rather than for soluble and TMD-containing non-secretome transcripts (Figure S2D).

### Gene ontology (GO) analysis

To determine those gene categories that are overrepresented in the population of SECReTE-containing genes, gene ontology (GO) enrichment analysis was conducted. When SECReTE10-positive genes were searched for GO enrichment (using all yeast genes as a background), unsurprisingly, membrane proteins were found to have a high enrichment score (fold enrichment = 67) (Figure 3A). The most SECReTE-enriched gene category was that comprising cell wall proteins (fold enrichment = 1.8) (Figure 3A). When 15 *NNY* repeats served as a threshold, the fold-change enrichment of the cell wall protein category increased to 4.8-fold (Figure 3B). To further characterize the mRNAs enriched with SECReTE, we divided the secretome and non-secretome into subgroups and calculated the fraction of transcripts containing SECReTE10 in each category. In agreement with the GO analysis, more than 90% of mRNAs coding for cell wall proteins possess SECReTE10 elements and the cell wall proteins were the most SECReTE-rich (Figure 3C). We found that 86% of mRNAs of proteins encoding both TMD and signal-sequence (SS) regions, as well as 84% of TMD-encoding secretome mRNAs, contain SECReTE10 (Figure 3C). Of these, mRNAs encoding tail-anchored (TA) proteins contain the lowest number of transcripts with SECReTE10 in the secretome (Figure 3C). TA proteins are known to translocate to the ER through an alternative pathway (GET) after being translated in the cytosol^40-42^, and their transcripts are not enriched on ER membranes ^22,23^. This could imply that SECReTE is more abundant in mRNAs undergoing translation on the ER. In contrast, transcript for non-secretome proteins (*i.e.* mitochondrial and cytonuclear) have the lowest abundance of SECReTE elements (Figure 3C).

**Figure 3.**
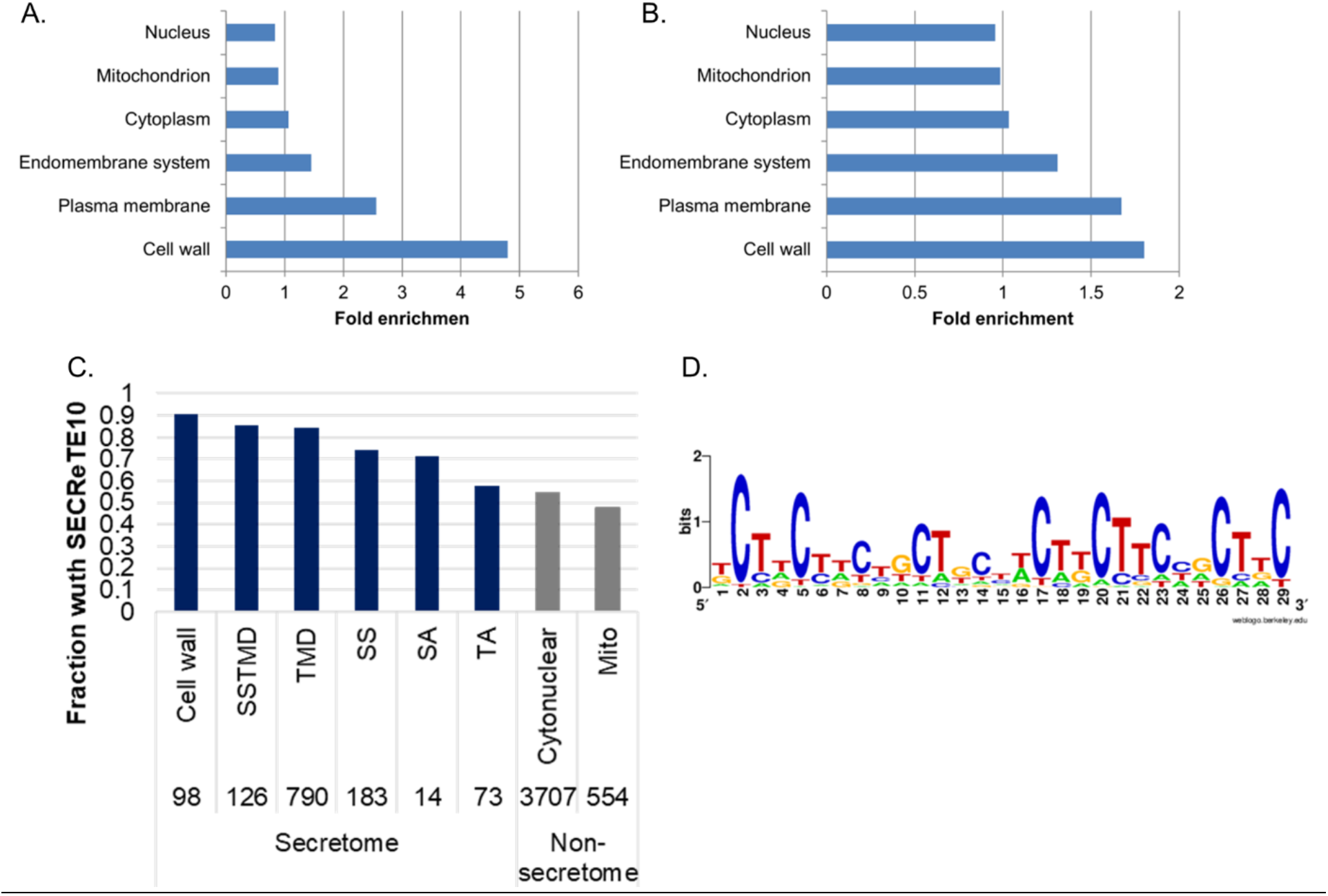
Cell wall proteins are highly enriched with SECReTE. **(A) GO annotation analysis for genes containing SECReTE10.** Genes encoding cell wall proteins, as well as membrane proteins, show the highest and most significant enrichment score. **(B) GO annotation analysis for genes containing SECReTE15.** Genes encoding cell wall proteins are the most enriched with SECReTE. **(C) SECReTE10 abundance in different groups of genes.** More than 90% of mRNAs encoding proteins annotated to localize to the cell wall contain SECReTE. High SECReTE abundance was also noticed in other secretome groups except tail-anchored (TA) proteins. Mitochondrial mRNAs (Mito) have low SECReTE abundance. Numbers above bars represent the number of genes in each group. **(D) MEME analysis of cell wall transcripts.** A motif similar to SECReTE was revealed in cell wall transcripts using MEME. Numbers on the x axis indicate base number.

Since SECReTE is highly enriched in mRNAs coding for cell wall proteins, we wanted to know if it could be discovered using an unbiased motif search tool. For that, we analyzed the mRNA sequences of cell wall proteins using MEME to identify mRNA motifs. The most significant result obtained highly resembled the SECReTE10 repeat with either U or C (Figure 3D). Importantly, we did not detect a protein motif within this mRNA motif, eliminating the possibility that the SECReTE element is dependent on the protein sequence.

### SECReTE enrichment in mSMPs is found in both prokaryotes and higher eukaryotes

Conservation or convergence in evolution are strong indications of significance. To check whether SECReTE enrichment in mSMPs is found in higher and lower organisms (*e.g.* humans and *B. subtilis*) we analyzed these genomes. In humans, as in *S. cerevisiae*, SECReTE10 gave the most significant separation between RNAs encoding secretome and non-secretome proteins, based on ROC analysis (Figure 4A). After verifying that SECReTE10 does not correlate with gene length, 10 *NNY* repeats served as a threshold to define presence of the SECReTE motif. As in yeast, SECReTE is enriched in the second and third codon positions of secretome transcripts, in comparison to non-secretome transcripts (Figure 4B). Also, a larger fraction of secretome transcripts that lack TMDs contain SECReTE, as compared to non-secretome transcripts bearing TMDs (Figure 4C). Interestingly, transcripts encoding glycophosphatidylinositol (GPI)-anchored proteins, which are equivalent to cell wall proteins, were found to be highly enriched with SECReTE. In contrast, tail-anchored genes, as well as mitochondrial and cytonuclear genes, have a low SECReTE abundance as seen in yeast (Figure 4D). We also detected a high abundance of SECReTE10 in genes encoding secretome proteins from *B. subtilis*, in comparison to those encoding non-secretome proteins (Figure 4E).

**Figure 4.**
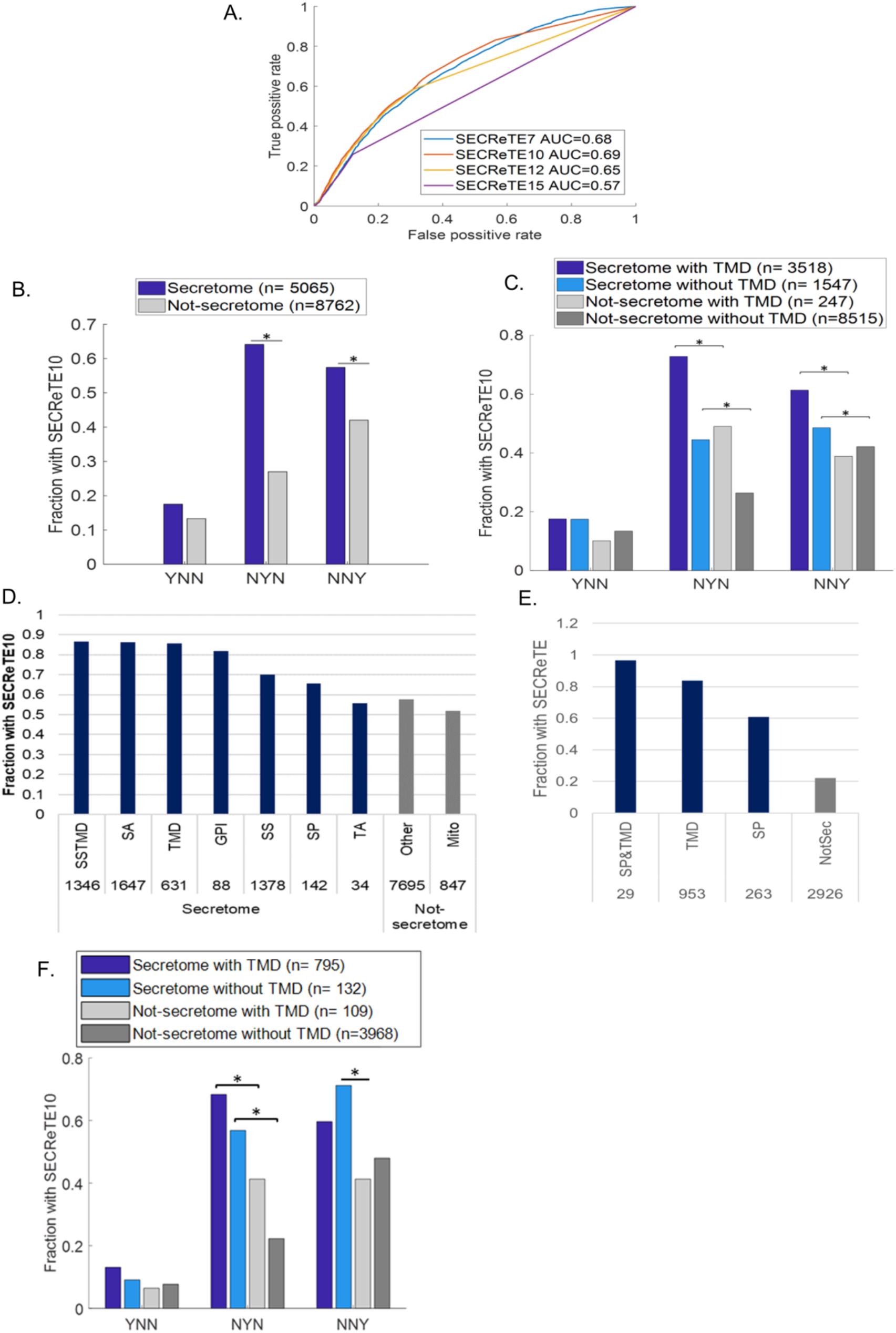
SECReTE is found in the human genome. **(A)** SECReTE10 maximizes the ability to classify secretome genes in human. ROC curves were plotted for each of the indicated thresholds. Secretome genes were used as the true positive set and non-secretome genes as the true negative set. The AUC (area under the curve) of SECReTE10 was the highest. **B. SECReTE is highly abundant in the mRNAs of human secretome proteins.** SECReTE10 abundance was calculated for each codon position separately. SECReTE abundance in human mSMPs is most significant in the second position of the codon, but highly significant differences were also detected in the third position. \**p*≤ 3.73E-68 **C. SECReTE is highly abundant in mRNAs coding for soluble secretome proteins in humans.** SECReTE10 presence was examined separately for TMD-containing proteins and soluble secreted proteins. A higher fraction of mRNAs coding for soluble secreted proteins (Secretome without TMD; cyan) contains SECReTE in comparison to non-secretome transcripts without a TMD (light gray). The fraction of soluble secreted proteins having SECReTE in the third position is larger than that of TMD-containing non-secretome proteins (NNY) and is significant. n represent the number of genes in each group. * *p≤* 3.49E-12. **(D) SECReTE10 abundance in different groups of genes.** High SECReTE abundance was observed for other secretome protein groups, except tail-anchored (TA) proteins. Mitochondrial mRNAs (Mito) have low SECReTE abundance. Numbers above bars represent the number of genes in each group. **(E) SECReTE10 abundance in *B. subtilis***. SECReTE10 abundance was scored and was observed to be higher in mRNA coding for genes encoding secretome proteins (*i.e.* SS&TMD, TMD, and SS) as compared to those encoding non-secretome (Non-Sec) proteins. Numbers under bars represent the number of genes in each group. **(F) SECReTE10 abundance in *S. pombe*** SECReTE10 abundance was calculated for each codon position separately for TMD-containing proteins and soluble secreted proteins. A higher fraction of mRNAs coding for soluble secreted proteins (Secretome without TMD; cyan) contains SECReTE in comparison to non-secretome transcripts, either with or without a TMD (Non-secretome with TMD; dark gray, Non-secretome without TMD; light gray). The fraction of soluble secreted proteins having SECReTE in the third position is larger than that of TMD-containing non-secretome proteins (NNY) and is significant. n represent the number of genes in each group. n represent the number of genes in each group. * *p≤* 5.63E-3.

### Analysis of the *Schizosaccharomyces pombe* genome suggests that SECReTE may arise via convergent evolution

We next asked if SECReTE is evolutionarily conserved via inheritance or had it emerged independently through convergent evolution. To differentiate between conservation and convergence we analyzed the genome of the fission yeast, *S. pombe*, for the presence and position of SECReTE in secretome and non-secretome transcripts. As found for *S. cerevisiae*, SECReTE is enriched (in the second and third codon positions) in a larger fraction of *S. pombe* mSMPs that lack TMDs, as compared those containing TMDs or to non-secretome transcripts that either bear or lack TMDs (Figure 4F). Next, we aligned orthologous genes encoding secretome proteins from *S. cerevisiae* to those of *S. pombe* (457 genes total), and examined whether SECReTE is found in the same position within the gene. However, we found that only a minor portion of SECReTE motifs (66) were shared spatially between orthologous genes. This finding implies that SECReTE motifs are more likely to have arisen via convergence than by conservation.

### Mutations in SECReTE affect the secretion of endogenous secretome proteins

To further understand the significance of SECReTE and validate its importance to yeast cell physiology, we examined its relevance by elevating or decreasing the signal in selected genes. Three representative genes were chosen, based on their relatively short gene length, a detectable phenotype upon their deletion, and their function in different physiological pathways. These genes included: *SUC2*, which encodes a soluble secreted periplasmic enzyme; *HSP150*, which encodes a soluble media protein; and *CCW12*, which encodes a GPI-anchored cell wall protein. The overall SECReTE signal of the genes was increased by substituting any A or G found in the third codon position with a T or C, respectively, thereby enriching SECReTE presence along the entire gene [(+)SECReTE]. The reverse substitution, converting T to A or C to G, decreased the overall SECReTE signal [(-)SECReTE]. The number of motifs present in each gene before and after SECReTE addition/reduction is shown in Table S3. Crucially, these modifications were designed to ensure that only the SECReTE attribute of the mRNA sequence was altered, while no alterations in the encoded amino acid sequence were made. Furthermore, changes in the stability of the mRNA secondary structure (free energy) and the Codon Adaptation Index (CAI)^40^ were kept to within a similar range (Table S3). SECReTE mutations in *SUC2, HSP150*, and *CCW12* are shown along the length of the gene, using a minimum threshold of either 1 NNY repeats or 10 NNY repeats, as shown in Figure S3 (A-C; upper and lower parts, respectively).

### SECReTE mutations in *SUC2* alter invertase secretion

*SUC2* codes for different forms of invertase translated from two distinct mRNAs, short and long, which differ only at their 5’ ends. While the longer mRNA codes for a secreted protein that contains a signal sequence, the signal sequence is omitted from the short isoform, which codes for a cytoplasmic protein. Secreted Suc2 expression is subjected to glucose repression; however, under inducing conditions (*i.e.*, glucose depletion), Suc2 is trafficked through the secretory pathway to the periplasmic space of the cell. There, it catalyzes the hydrolysis of sucrose to glucose and fructose, this enzymatic activity being responsible for the ability of yeast to utilize sucrose as a carbon source and can be measured by a biochemical assay (*i.e.* invertase activity), both inside and outside of the cell. The effect of SECReTE mutations on Suc2 function was tested by examining the ability of mutants to grow on sucrose-containing media by drop-test. Interestingly, the growth rate of *SUC2*(-)SECReTE on sucrose plates was decreased, while the *SUC2*(+)SECReTE mutant exhibited better growth in comparison to WT cells (Figure 5A), even though no growth change was detected on YPD plates. These findings suggest that SECReTE strength affects the secretion of Suc2. These changes in Suc2 secretion could result from changes in *SUC2* transcription, Suc2 production, and/or altered rates of secretion. To distinguish between possibilities, WT cells, *suc2*Δ, and *SUC2* SECReTE mutants were subjected to invertase assays. The invertase assay enables the quantification of secreted Suc2, as well as internal Suc2, by calculating the amount of glucose produced from sucrose. As expected, under glucose repressing conditions (*e.g.* 2% glucose) the levels of both secreted and internal Suc2 were very low. When cells were grown on media containing low glucose (*e.g.* 0.05% glucose) to promote the expression of the secreted enzyme, secreted Suc2 levels were altered due to changes in SECReTE. Corresponding to the drop-test results, a significant decrease in secreted invertase was detected with *SUC2*(-)SECReTE cells, while a significant increase was detected with *SUC2*(+)SECReTE cells, in comparison to WT cells. No Suc2 secretion was detected from *suc2*Δ cells (Figure 5B, secreted). If SECReTE mutations affect the efficiency of Suc2 secretion, but not its synthesis, then Suc2 accumulation would be expected to occur in *SUC2*(-)SECReTE cells. Likewise, as a decrease of internal invertase would be expected to occur in *SUC2*(+)SECReTE cells. However, this was not the case as the internal amount of Suc2 was decreased in *SUC2*(-)SECReTE cells and was slightly increased in *SUC2*(+)SECReTE cells (Figure 5B, internal). These findings suggest that SECReTE alterations in *SUC2* likely affect protein production.

**Figure 5.**
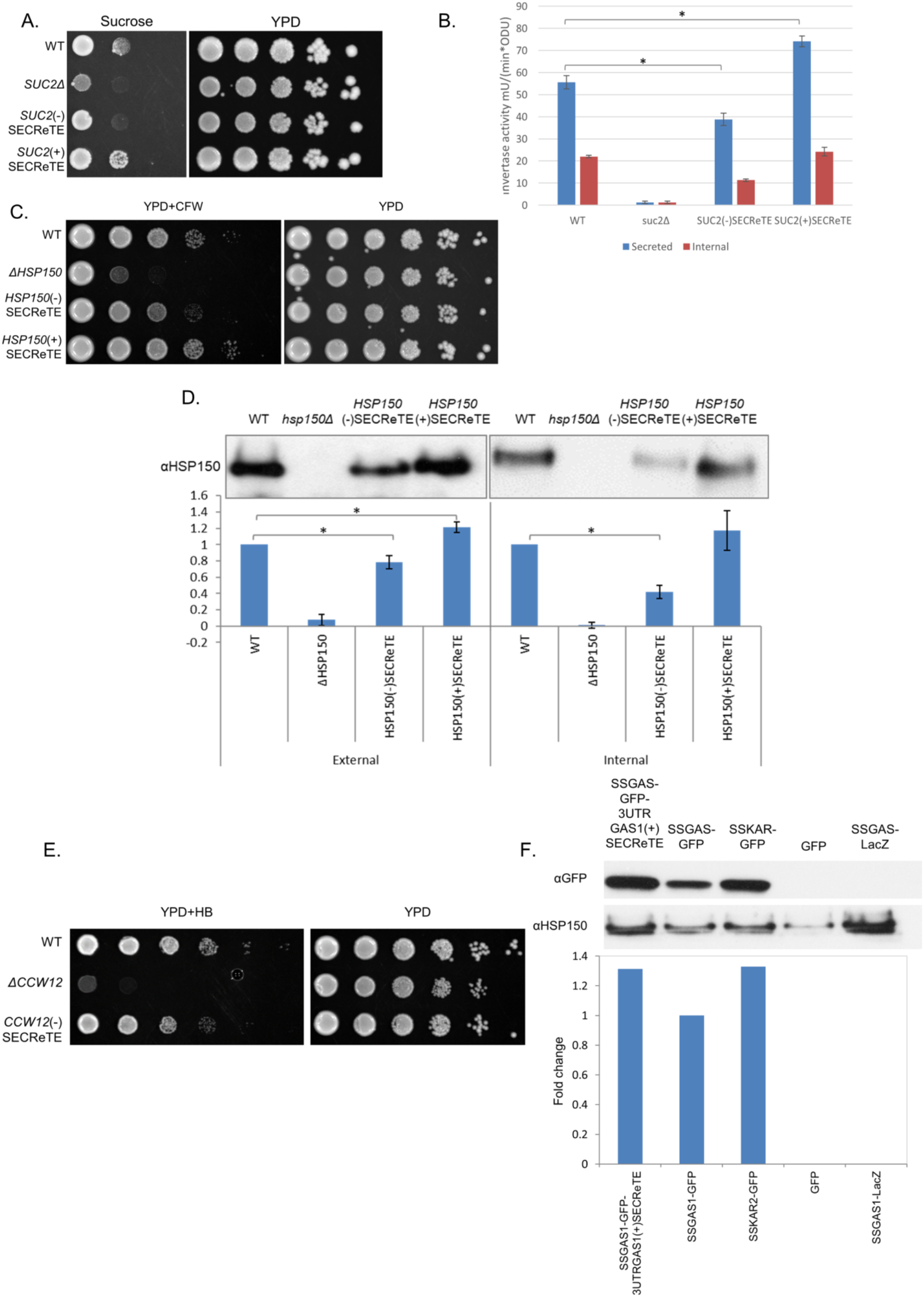
The levels of secretion of endogenous and exogenous proteins are affected by SECReTE strength. **(A) SECReTE enhances the ability to grow on sucrose.** The ability of WT, *suc2*Δ, *SUC2*(+)SECReTE and *SUC2*(-)SECReTE yeast to grow on sucrose was examined by drop-test. Cells were grown to mid-log on glucose-containing YPD medium, prior to serial dilution and plating onto sucrose-containing synthetic medium or YPD. Cells were grown for 2 days prior to photodocumentation. The *SUC2*(-)SECReTE mutant exhibited reduced growth than WT cells, while *SUC2*(+)SECReTE cells exhibited better growth. *suc2*Δ cells were unable to grow on sucrose-containing medium. **(B) SECReTE enhances invertase secretion.** The indicated strains from *A* were subjected to the invertase secretion assay. Both internal and secreted invertase activity was measured in units after glucose de-repression. Both activities were reduced in *SUC2*(-)SECReTE cells and elevated in *SUC2*(+)SECReTE cells. Error bars represent the standard deviation from three experimental repeats. \**p*≤0.05. **(C) SECReTE enhances the ability to grow on calcofluor white.** The ability of WT, *hsp150*Δ, *HSP150*(+)SECReTE and *HSP150*(-)SECReTE cells to grow on CFW was examined by drop-test. Cells were grown to mid-log on YPD, prior to serial dilution and plating on YPD alone or YPD plates containing CFW, and incubated at 30°C. Cells were grown for 2 days prior to photodocumentation. The *HSP150*(-)SECReTE mutant exhibited hypersensitivity in comparison to WT cells, while *HSP150*(+)SECReTE cells were less sensitive. *hsp150*Δ cells grew poorly on medium containing CFW. **(D) SECReTE enhances Hsp150 secretion.** The indicated strains from *C* were subjected to the Hsp150 secretion assay. Cells were grown to mid-log phase at 37°C for 4hrs and examination in cell lysates (internal) or medium (external) by Western analysis using anti-Hsp150 antibodies. External Hsp150 was decreased in *HSP150*(-)SECReTE cells in comparison to WT, while it was increased in the *HSP150*(+)SECReTE strain. Internal Hsp150 was decreased in *HSP150*(-)SECReTE cells and also slightly in *HSP150*(+)SECReTE cells, in comparison with WT cells. No internal nor external Hsp150 was detected in the lysate or medium derived from *hsp150*Δ cells, respectively. Band intensity was quantified using ImageJ and presented in the histogram below. The graphs represent the ratio of the intensity of all samples relative to that of WT. **(E) SECReTE enhances the ability to grow on hygromycin B.** The ability of WT, *ccw12*Δ, and *CCW12*(-)SECReTE cells to grow on HB was examined by drop-test. Cells were grown to midlog on glucose-containing YPD medium, prior to serial dilution and plating onto HB-containing YPD or YPD alone. Cells were grown for 2 days prior to photodocumentation. The *CCW12*(-)SECReTE strain was more sensitive to HB stress in comparison to WT cells. *ccw12*Δ cells were unable to grow on medium containing HB. **(F) SECReTE enhances secretion of an exogenous protein, SSGAS1-GFP.** Yeast expressing SSGAS1 -GFP3’UTRGAS 1(+)SECReTE, SSGAS1-GFP, SSKAR2-GFP, GFP, and SSGAS1-LacZ from plasmids were grown to mid-log phase on synthetic medium containing 2% raffinose and shifted to 3% galactose-containing medium for 4hrs. Proteins expressed from the different strains were TCA precipitated from the medium and the precipitates resolved by SDS-PAGE. GFP was detected with an anti-GFP antibody, while Hsp150 was detected with an anti-Hsp150 antibody and was used as a loading control. Band intensity was quantified using ImageJ; intensity was scored relative to SSGAS1-GFP secretion. Addition of the *GAS1* 3’UTR mutated to contain SECReTE improved the secretion of SS-Gas1 and was comparable to that of SSKAR2-GFP. GFP lacking a SS was not secreted and SSGAS1-LacZ was used as a negative control.

### SECReTE mutations alter Hsp150 secretion and cell wall stability

Next, we wanted to study the importance of SECReTE in *HSP150*. Hsp150 is a component of the outer cell wall and while the exact function of Hsp150 is unknown, it is required for cell wall stability and resistance to cell wall-perturbing agents, such as Calcofluor White (CFW) and Congo Red (CR). While *hsp150*Δ cells are more sensitive to cell wall stress, the overproduction of Hsp150 increases cell wall integrity^43^. Hsp150 is secreted efficiently into the growth media and its expression is increased upon heat shock^44,45^. The effect of modifying SECReTE in *HSP150* was examined via drop-test by testing the sensitivity of *HSP150*(-)SECReTE and *HSP150*(+)SECReTE cells to added CFW, in comparison to WT and *hsp150*Δ cells. As can be seen from Figure 5C, while the *HSP150*(-)SECReTE strain was more sensitive to CFW as compared to WT cells, the *HSP150*(+)SECReTE strain was more resistant to CFW. As expected, *hsp150*Δ cells are the most susceptible to CFW (Figure 5C). *HSP150* strains were also subjected to Western blot analysis to measure levels of the mutant proteins. Since *HSP150* secretion is elevated upon heat-shock^44,45^, cells were grown at 37°C before protein extraction. Protein was extracted from both the growth medium and cells to detect both external and internal protein levels, respectively. The amount of Hsp150 secreted to the medium was decreased in *HSP150*(-)SECReTE cells and elevated in *HSP150*(+)SECReTE cells, in comparison to WT cells (Figure 5D). Similar to Suc2, the internal amount of Hsp150 was also decreased in *HSP150*(-)SECReTE cells, as compared to WT cells (Figure 5D). This could mean that secretion *per se* was not significantly attenuated by the reduction in SECReTE strength. As the internal amount of Hsp150 in *HSP150*(+)SECReTE cells was similar to that of WT cells, we concluded that SECReTE alteration in *HSP150* also likely affects protein production.

### SECReTE mutations in *CCW12* alter cell wall stability

*CCW12* encodes a GPI-anchored cell wall protein that localizes to regions of the newly synthesized cell wall and maintains wall stability during bud emergence and shmoo formation. Deletion of *CCW12* was shown to cause hypersensitivity to cell wall destabilizing agents, like hygromycin B (HB)^46,47^. Since the SECReTE score is very high in *CCW12*, it was not possible to further increase the signal. Therefore, we generated only *CCW12*(-)SECReTE cells and tested their ability to grow on HB-containing plates. As seen with *HSP150*(-)SECReTE (Figure 5C), we found that the *CCW12*(-SECReTE mutation rendered cells sensitive to cell wall perturbation, in comparison to WT cells (Figure 5E).

### SECReTE addition affects secretion of an exogenous naïve protein

The ability of SECReTE addition to improve the secretion of an exogenous protein would not only be substantial evidence for its importance, but also could be a useful, low-cost, industrial tool to improve the secretion of recombinant proteins without changing protein sequence. To test that, we employed a *GFP* transcript construct bearing the encoded SS of Gas1 (SSGas1) at the 5’ end. SSGasI enables the secretion of GFP protein to the medium, although its secretion was not as efficient in comparison to other SS-fused GFP proteins, such as SSKar2 (Figure 5F). To potentially improve the secretion of SSGas1, we added an altered 3’UTR sequence of Gas1 that contained SECReTE [*i.e.* in which all A’s and G’s were replaced with T’s and C’s, respectively; SSGasI-3’UTRGASI(+)SECReTE]. We then tested the effect of SECReTE addition upon the secretion of GFP into the media. We found that the addition of SECReTE to the 3’UTR of GasI-GFP improved the secretion of GFP secretion into the media, in comparison to SSGasI-GFP, and was similar to that of SSKar2-GFP (Figure 5F).

### The effect of SECReTE mutations on mRNA levels

As protein levels may be altered by (-)SECReTE and (+)SECReTE mutations (Figure 5B, D, and F), we examined whether changes in gene transcription or mRNA stability are involved. Quantitative real-time (qRT) PCR was employed to check whether mRNA levels of *SUC2, HSP150*, and *CCW12* are affected by SECReTE strength. We found that *SUC2*(-)SECReTE mRNA levels were almost 30% lower than in *SUC2* WT cells, while *SUC2*(+)SECReTE levels were ~50% higher than WT (Figure S4A). This change in mRNA levels might be the cause for the ability of *SUC2*(+)SECReTE mutant to increase protein production and, therefore, grow better on sucrose-containing medium (Figure 5A,B).

The effect of SECReTE mutation on *HSP150* mRNA levels was also studied. Interestingly, we found that the mRNA level of *HSP150*(-)SECReTE was similar to WT, while that of *HSP150*(+)SECReTE was slightly decreased (Figure S4B). Thus, the change in Hsp150 protein levels and sensitivity to CFW due to SECReTE alteration (Figure 5C and D) is not explained by changes in mRNA levels. Likewise, SECReTE mutations in *CCW12*(-)SECReTE did not cause a significant change in its mRNA level (Figure S4C). Therefore, the increased sensitivity of *CCW12*(-) SECReTE to HB (Figure 5E) is not due to a decrease in *CCW12* mRNA.

### The effect of SECReTE mutation on *SUC2* mRNA localization

To test whether SECReTE has a role in dictating mRNA localization, we visualized *SUC2* mRNA by single-molecule FISH (smFISH) using specific fluorescent probes and tested the influence of SECReTE alteration on the level of *SUC2* mRNA co-localization with ER. We used Sec63-GFP as an ER marker and calculated the percentage of granules per cell that co-localized either with or not with the ER, or were adjacent to the ER. The level of co-localization between *SUC2*(-)SECReTE mRNA granules and Sec63-GFP was found to decrease slightly in comparison to WT *SUC2* mRNA granules (Figure 6A and B). In contrast, we observed a significant increase of ~50% in the level of co-localization of *SUC2*(+)SECReTE mRNA granules with the ER, in comparison to WT *SUC2* mRNA (Figure 6A and B). These findings suggest that SECReTE has role in the targeting of *SUC2* mRNA to the ER.

**Figure 6.**
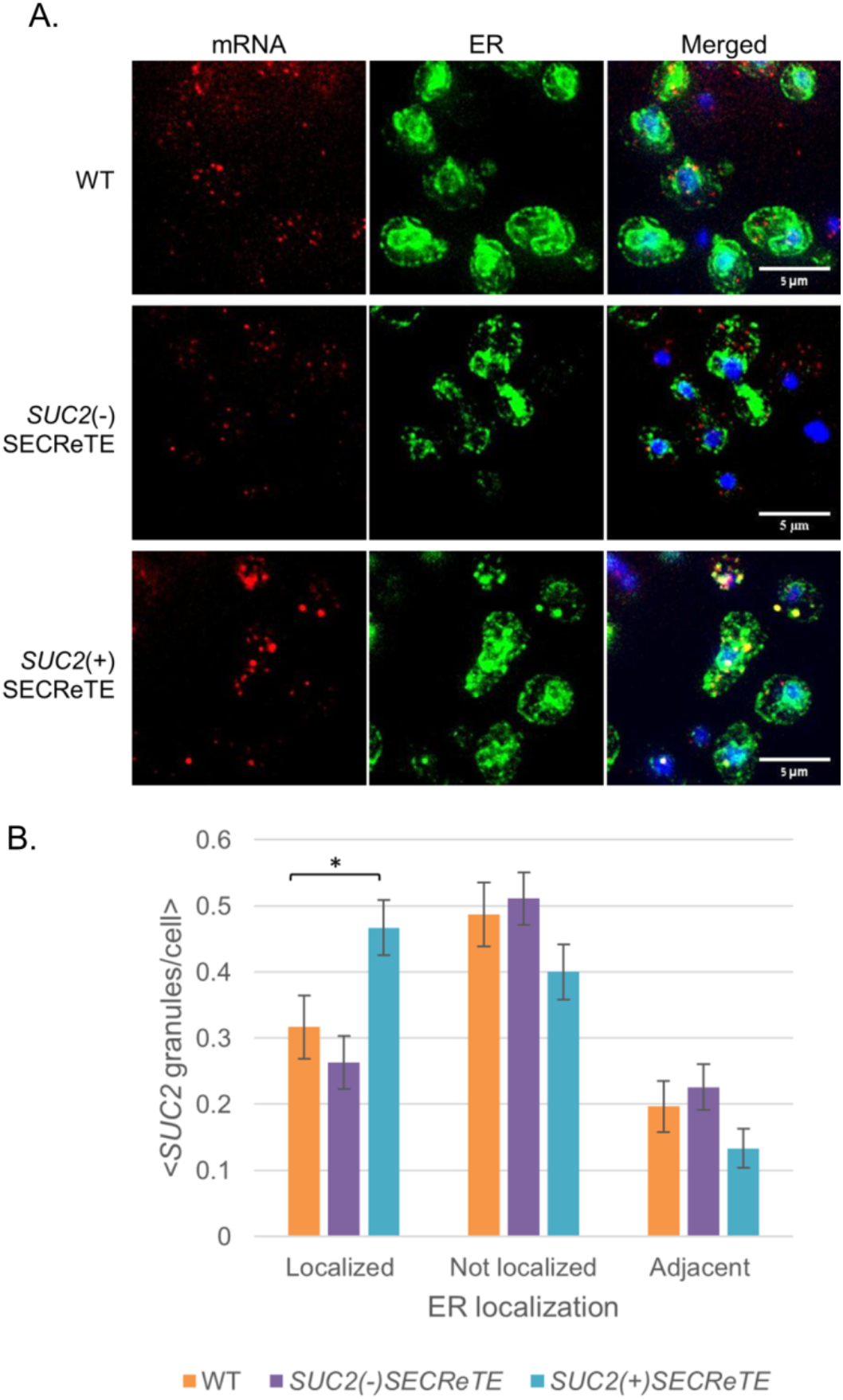
SECReTE enhances *SUC2* mRNA localization to the ER. **(A) Visualization of endogenously expressed *SUC2*(+)SECReTE and *SUC2*(-)SECReTE mRNAs using smFISH.** Yeast endogenously expressing WT *SUC2*, *SUC2*(+)SECReTE, or *SUC2*(-)SECReTE and Sec63-GFP from a plasmid were grown to mid-log phase on SC medium containing 2% glucose prior to shifting cells to low glucose-containing medium (0.05% glucose) to induce *SUC2* expression. Cells were processed for smFISH labeling using non-overlapping, TAMRA-labeled, FISH probes complementary to *SUC2*. **B. Quantification of *SUC2*(+)SECReTE and *SUC2*(-)SECReTE mRNA localization to the ER.** The percentage of granules that are co-localized, not co-localized, or adjacent to Sec63-GFP labeled ER was scored in each cell. The histogram shows the average score for at least ~60 cells and ~250 *SUC2* granules for each strain. \**p* =0.019.

### Identification of potential SECReTE-binding proteins

To further elucidate the role of SECReTE it is essential to identify its binding partners, presumably RBPs. Large-scale approaches were previously used to identify mRNAs that are bound >40 known RBPs in yeast^48-50^. To obtain a list of potential SECReTE-binding proteins (SBPs) we searched the datasets for RBPs that bind mRNAs highly enriched with SECReTE. For each RBP, we calculated what fraction of its bound transcripts contain SECReTE10. RBPs found to bind large fractions of SECReTE10-containing mRNAs included Bfr1, Whi3, Puf1, Puf2, Scp160, and Khd1 (Figure 7A), and were all previously shown to bind mSMPs^48-50^. To test which of these candidates bind SECReTE, each of the genes these RBPs was deleted in either WT or *HSP150*(+)SECReTE cells. We hypothesized that the deletion of a genuine SBP might confer hypersensitivity to CFW and eliminate the growth rate differences between WT and *HSP150*(+)SECReTE cells observed on CFW-containing plates (Figure 5C). When *PUF1, PUF2*, or *SHE2* were deleted we found that *HSP150*(+)SECReTE strain was still more resistant to CFW than WT cells (Figure S5). One possible explanation for this lack of effect is that these RBPs either do not bind *HSP150* or that they are redundant with other SBPs. However, we did find that the deletion of either *WHI3* or *KHD1* eliminated the differences between WT and *HSP150*(+)SECReTE strains on CFW-containing plates (Figure 7B). This suggests Whi3 and Khd1 bind *HSP150* mRNA and possibly other secretome mRNAs, and even WT cells alone were rendered more sensitive to CFW in their absence (Figure 7B).

**Figure 7.**
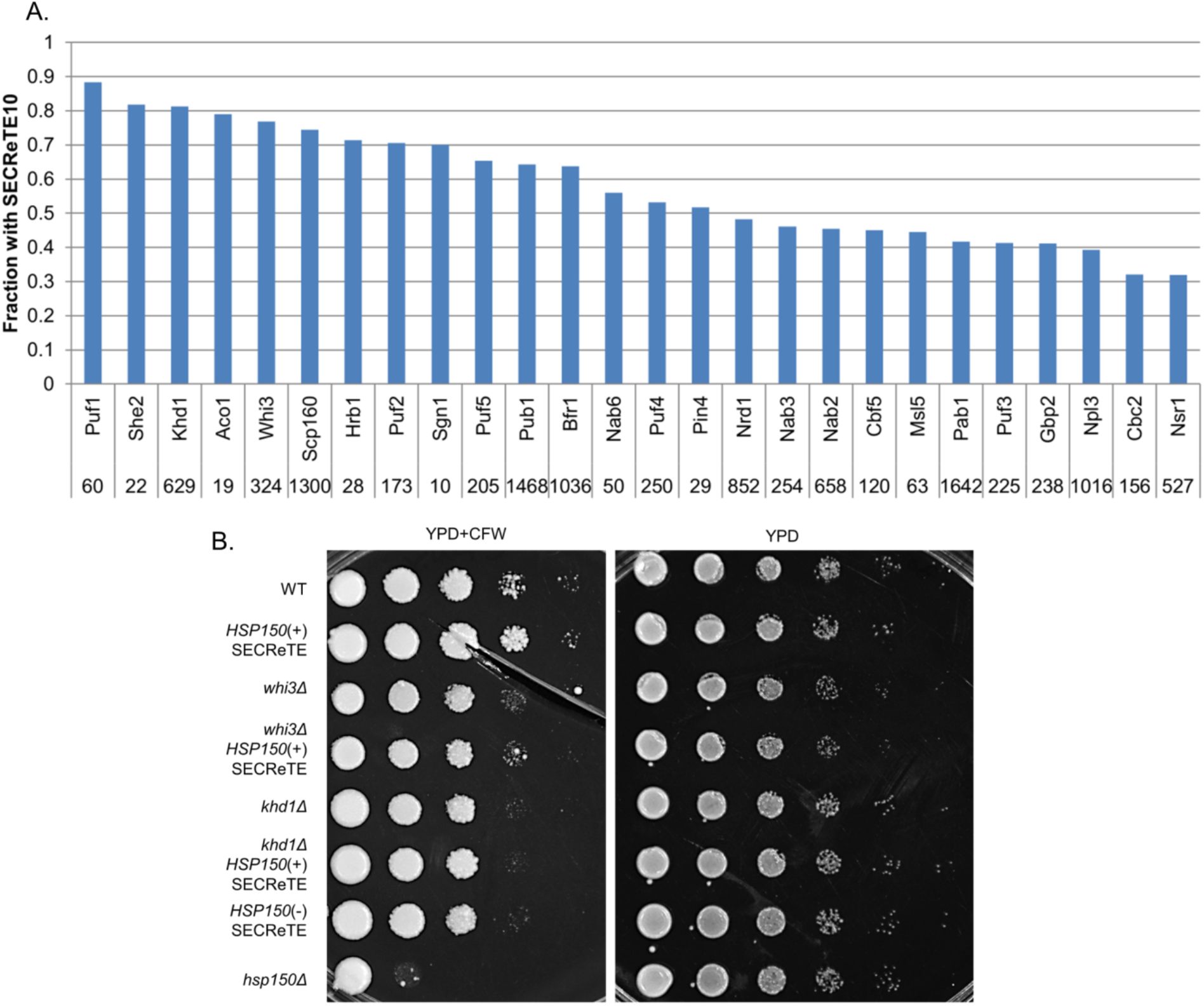
Identification of potential SECReTE-binding proteins. **(A) Identification of SECReTE10-containing transcripts in RNA-binding protein pulldown studies.** The number and fraction of SECReTE10-containing mRNAs from the total mRNAs bound to the indicated RBPs is shown. The microarray analysis data used to generate the histogram was published in ^49-51^. **(B) Identification of potential SECReTE-binding partners.** WT cells and either WT or *HSP150*(+)SECReTE cells deleted for genes encoding the indicated RBPs (*e.g.* Whi3, and Khd1) were grown to mid-log phase on YPD at 30°C, prior to serial dilution and plating onto either solid YPD medium or YPD containing CFW. Yeast were grown 2 days prior to photodocumentation.

## Discussion

The sorting of proteins to their proper destination is crucial for cellular organization and normal function. While the information for protein localization can reside within the protein sequence (*e.g.* protein targeting sequences), the spatial localization of an mRNA may also be important for protein proper targeting cell^1,2^. For example, mSMPs localize to the surface of the ER independently of translation and that localization requires elements within the transcript that are presumably recognized by an ER-localized RBP (see reviews^8,9,51^). It was shown previously that ER-targeted TMD-containing proteins are highly enriched with amino acids containing uracil-rich codons^30^ and, thus, their ORFs are enriched with pyrimidines^26^. Nevertheless, mRNAs coding for secretome proteins that do not contain TMDs were also found to be enriched on ER membranes ^2,H52^. Therefore, an additional mechanism or element appears necessary to confer mSMP localization. Here, we identify features that characterize all mSMPs, either encoding a TMD or not, and discovered a repetitive motif consisting of >10 consecutive *NNY* repeats. This motif, termed SECReTE, is not restricted to transcripts coding for TMD-containing proteins, but can be found in higher abundance in all secretome transcripts, from prokaryotes (*e.g. B. subtilis*) to yeast (*S. cerevisiae* and *S. pombe*) to humans (Figures 1,4). By analyzing the *S. pombe* genome it was discovered that SECReTE tends to positioned differently in orthologous genes encoding secretome proteins, in comparison to those in *S. cerevisiae*. This implies that SECReTE enrichment in mSMPs may evolved independently and, therefore, is convergent in evolution. Importantly, this convergence further emphasizes the significance and functionality of the SECReTE motif itself.

To better characterize SECReTE, we first determined the number of *NNY* repeats that can serve as a threshold to verify its presence and found that ten (*i.e.* SECReTE10) were found to constitute a genuine motif, rather than a random occurrence and enabled significant separation between secretome and non-secretome mRNAs (Figure 1). SECReTE abundance was calculated separately for each position of the codon and while being barely present in the first position (Figure 2A, *YNN*), it was highly represented in the second and third positions in mSMPs (*NYN* and *NNY*, respectively), in comparison to non-mSMPs. Interestingly, the SECReTE10-containing fraction of transcripts coding for soluble secreted proteins is larger than that of mRNAs encoding secreted membrane proteins, suggesting that SECReTE enrichment is not merely due to the high fraction of TMD-containing genes in the secretome (Figure 2B). Importantly, when encoded TMD sequences were removed from the analysis, SECReTE10 was found to be more abundant in the third position of the codon in secretome transcripts (Figure 2C).By analyzing the datasets of both Jan *et al.*^22^ and Chartron *et al*^23^, we verified that a higher fraction of SECReTE10-containing transcripts is enriched on ER-bound ribosomes (Figure S1A) and in polysomes extracted from the membrane fraction (Figure S1B), as well as in the membrane fraction itself (Figure S1C). In contrast, transcripts with SECReTE10 were not enriched on mitochondria-bound ribosomes (Figure S1D). Permutation analysis confirmed that SECReTE enrichment in mSMPs is not arbitrary and demonstrated that it is independent of codon composition (Figure S2).

Although SECReTE10 enables the classification of mSMPs (Figures 1C and 4C), the separation between secretome and non-secretome is not absolute and mRNAs coding for non-secretome proteins also may contain SECReTE sequences. While this might suggest that the motif is not completely defined, it might also imply that SECReTE plays a role in non-secretome mRNAs, perhaps in ER localization. There is an ongoing debate regarding whether mRNAs encoding cytosolic proteins also localize to the ER and even be translated by ER-associated ribosomes^53,54^. The idea that ER can support the translation of both secretory and cytosolic proteins was initially proposed by Nicchitta and colleagues^14-16,51,55^. Furthermore, they suggested that since translation initiation can start before the emergence of the signal sequence, ER-bound ribosomes would not distinguish between mRNAs and, therefore, a certain amount of mRNAs encoding cytosolic proteins can be tethered to the ER membrane^16,23,51^. The fact that a large fraction of mRNAs encoding cytosolic proteins also contain SECReTE raises the possibility that their targeting to the ER is intentional and that this motif plays a role in it.

Gene ontology analysis revealed that genes encoding cell wall proteins are most enriched with SECReTE (Figure 5A-C). In contrast, TA-protein encoding transcripts are less-enriched with SECReTE (Figure 5C), perhaps since they are not enriched on ER membranes^22,23^ and their translation products only translocate to the ER after full translation in the cytosol^40-42^. This implies that SECReTE is more abundant in mRNAs meant to be translated on the ER. Importantly, the SECReTE motif was also found by an unbiased method for discovery using the MEME server to identify common motifs in cell wall genes. This result verifies our original identification of SECReTE and its importance is further enhanced by the findings that it is conserved/convergent with bacteria and humans (Figure 4). As in yeast, human mSMPs (*i.e.* codon regions) are more enriched with SECReTE than non-secretome transcripts, independently of TMD presence (Figure 4B-C). Unlike yeast, however, human transcripts contain larger portions of UTR sequences and further analysis is required to calculate SECReTE abundance in the UTR regions.

The physiological relevance of SECReTE was explored by altering its enrichment in three mSMPs: *SUC2, HSP150*, and *CCW12* (Figure 5). Although the amino acid sequences were not altered by mutation, the functionality of these genes was. *SUC2* SECReTE mutant exhibited altered growth rates on sucrose-containing medium in comparison to WT cells, *i.e.* reduced growth when motif strength was decreased and better growth when motif strength was elevated (Figure 5A). This result corresponded with either a decrease or increase in both invertase synthesis and secretion, respectively (Figure 5A,B). *HSP150* SECReTE mutants also behaved differently, *i.e. HSP150*(-)SECReTE cells exhibited higher sensitivity to CFW in comparison to WT cells, while *HSP150*(+)SECReTE cells were more resistant (Figure 5C). Similarly, *CCW12*(-)SECReTE cells exhibited hypersensitivity to HB (Figure 5E). These findings strengthen the notion that SECReTE plays an important biological role in regulating the amount of protein secreted from cells. This was verified using an exogenous substrate, SS(GAS1)-GFP, whose secretion was significantly enhanced upon addition of the Gas1 3’UTR containing the SECReTE motif (Figure 5F). Moreover, strengthening of SECReTE not only increased protein production and secretion, it also enhanced the localization of *SUC2* transcripts to the ER (Figure 6).

Although SECReTE is present throughout evolution, it is not a strict sequence-based motif since a wide variety of pyrimidine-rich sequences fit its demands. This variability might allow for the preferential binding of specific mSMPs (or non-secretome-encoding mRNAs that contain SECReTE elements) to different SBPs under different conditions, depending upon secretory needs of the cells. While it is generally assumed that mRNA localization is required for local translation and proper positioning of the translated protein, SBP binding may provide spatial and temporal regulation of mRNA stability and protein synthesis^56,57^. Accordingly, correct mRNA localization is not redundant to protein localization, but is another level of regulation that affects protein production. Supportive of this model is Puf3, an RBP that targets its associated mRNAs to the surface of the mitochondria^58^. In addition to its mitochondrial targeting role, Puf3 binding regulates the translational fate of mRNAs. Specifically, Puf3 binding leads to mRNA decay and repressed translation on high glucose, but becomes phosphorylated and promotes translation under low glucose conditions^59,60^. Interestingly, alterations in *SUC2* and *HSP150* SECReTE motifs also support this model, as mutations altered the amount of secreted protein, but not the ratio between secreted and non-secreted protein (Figure 5B,D). Thus, SECReTE strength affects protein production, but not necessarily the rate of secretion. Moreover, since Suc2 and Hsp150 both contain a signal peptide, SECReTE alteration may not necessarily affect protein targeting, but only mRNA targeting. Yet, if localizing mRNAs to the ER is important for conferring efficient translation, either through mRNA stabilization or the regulation of protein production, then SECReTE presence and strength is expected to fill a regulatory role. If SECReTE affects mRNA stabilization, this would explain why we observed a significant decrease in *SUC2*(-)SECReTE mRNA levels and an increase in *SUC2*(+)SECReTE mRNA levels, in comparison to WT cells (Figure S4A). While this was not the case for either *HSP150* or *CCW12* (Figure S4B-C), the levels of protein production were elevated for Hsp150. Thus, SECReTE presence can affect the localization, stability, and translation of mRNAs (Figures 5, 6, and S3).

If SECReTE is a *cis* regulatory element, the question is who are its *trans-acting* partners? Large-scale approaches have been used to identify mRNAs that interact with known RBPs in yeast^48-50^. These analyses enabled the identification of Bfr1, Whi3, Puf1, Puf2, Scp160, and Khd1 as potential SBPs, based upon their ability to interact with known SECReTE-containing transcripts (Figure 7A). As a means of verification, we first deleted individual RBPs and determined whether this alleviated the growth differences between WT and *HSP150*(+)SECReTE cells on CFW-containing medium, as might be expected upon the removal of a *bona fide* SBP. While the deletion of *PUF1, PUF2*, or *SHE2* did not alter the increased resistance of *HSP150*(+)SECReTE cells to CFW, those of *KHD1* and *WHI3* did (Figure 7D and S4). This suggests that they may be SBPs and several indications support the idea that Whi3 and Khd1 serve in this regard. For example, Whi3 possesses an RNA recognition motif and was already identified as preferentially binding mSMPs, including *HSP150*^48,61^. Whi3 also binds *CLN3* mRNA and is important for the efficient retention of Cln3 at the ER^62^, as well as to destabilize *CLN3* and other mRNA targets^61^. In addition, the *whi3* deletion mutant is sensitive to cell wall perturbing agents, such as CFW and congo-Red^48^, and is synthetic lethal with the deletion of *CCW12* in a synthetic genetic analysis screen^47^. Thus, Whi3 is an attractive candidate SBP. The same can be said for Khd1, which interacts with hundreds of transcripts including many mSMPs^49^. These transcripts include *CCW12*^49^ and, correspondingly, Khd1 plays a role in the cell wall integrity signaling pathway^63^ However, Khd1 is best known for its association with *ASH1* mRNA and is required for both its translational repression and efficient localization to the bud tip^64^. *ASH1* mRNA, as well as mRNAs encoding polarity and secretion factors (*e.g. SRO7*), are physically bound to cortical ER and both are delivered to the bud tip via the same mechanism involving She2, She3, and Myo4/She1^65,66^. Importantly, both *ASH1* and *SRO7* have SECReTE10 motifs (data not shown). Thus, Khd1 interactions with SECReTE-containing mRNAs might potentiate their targeting to the ER.

Here we reveal and characterize a novel *cis* element enriched in mSMPs that affects the secretion of both endogenous and heterologous proteins, probably through the regulation of protein production. Although the mechanism is not yet clear, SECReTE binding to ER-associated SBPs may enhance transcript interactions with the ER, mRNA stabilization, or both, with the result being either increased translation efficiency and/or number of mRNAs translated on ER-bound ribosomes (see model, Figure 8). Our model supports the idea that mRNA plays an active role in its own targeting and this does not necessarily contradict the importance of co-translational localization, but rather provides another level of regulation. Thus, we believe that SECReTE plays an important physiological role in the fine-tuning of cellular secretion.

**Figure 8.**
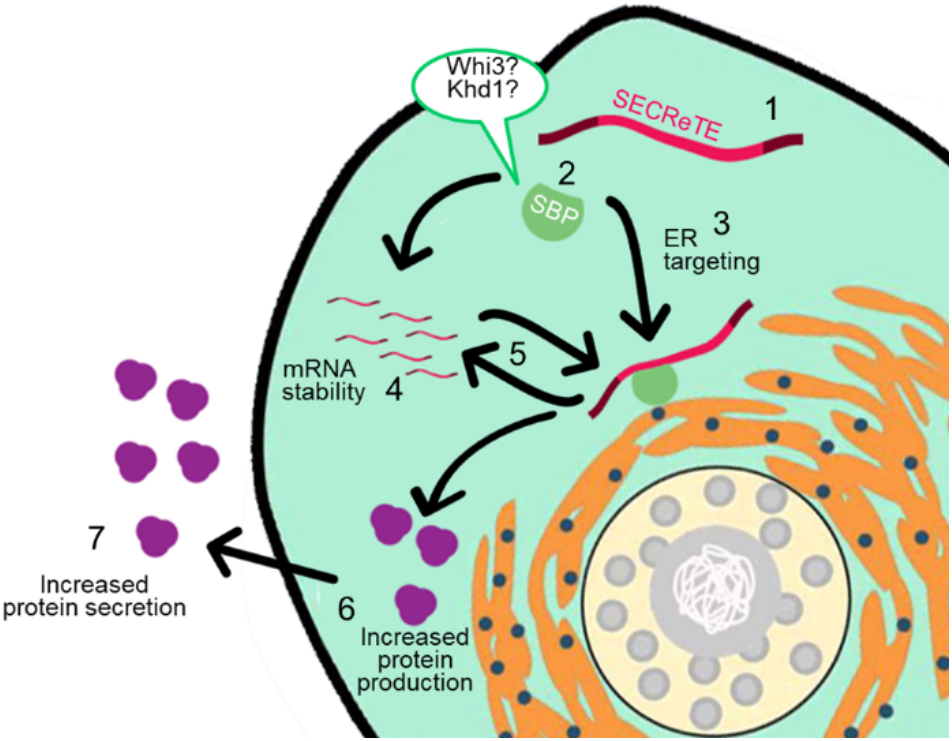
SECReTE plays an active role in protein secretion. SECReTE-containing transcripts (1) bind SBPs (2) and induce mRNA targeting to the ER (3), and/or confer mRNA stabilization (4). Targeting to the ER may provide spatial regulation and mRNA stabilization (5), leading to subsequent increases in protein production (6) and secretion (7).

Being both a unicellular and eukaryotic organism, *S. cerevisiae* is advantageous for the production of recombinant proteins as it grows quickly, is easy to culture, and secretes post-translationally modified proteins into the extracellular medium, which can facilitate their purification. Moreover, *S. cerevisiae* is a generally recognized as a safe (GRAS) organism, which makes it a favorable for use in the production of biopharmaceuticals^67,68^. Unfortunately, the natural capacity of *S. cerevisiae* secretory pathway is relatively limited and, thus, mechanisms that improve secreted protein production would be of significant benefit. Since SECReTE appears convergent in evolution, its use as an added RNA motif may prove to be a simple low-cost tool to improve recombinant protein production.

## Acknowledgements

The authors thank Jussi Jäntti and Maya Schuldiner for the gift of reagents, and Sivan Kaminski for data analysis. This study was supported by a grant to J.E.G from the Astrachan Olga Klein Fund, Weizmann Institute of Science. J.E.G. holds the Besen-Brender Chair in Microbiology and Parasitology, Weizmann Institute of Science. Y.P. acknowledges support from the European Research Council for the tRNAProlif grant.

## Supplementary Information

**Figure S1.**
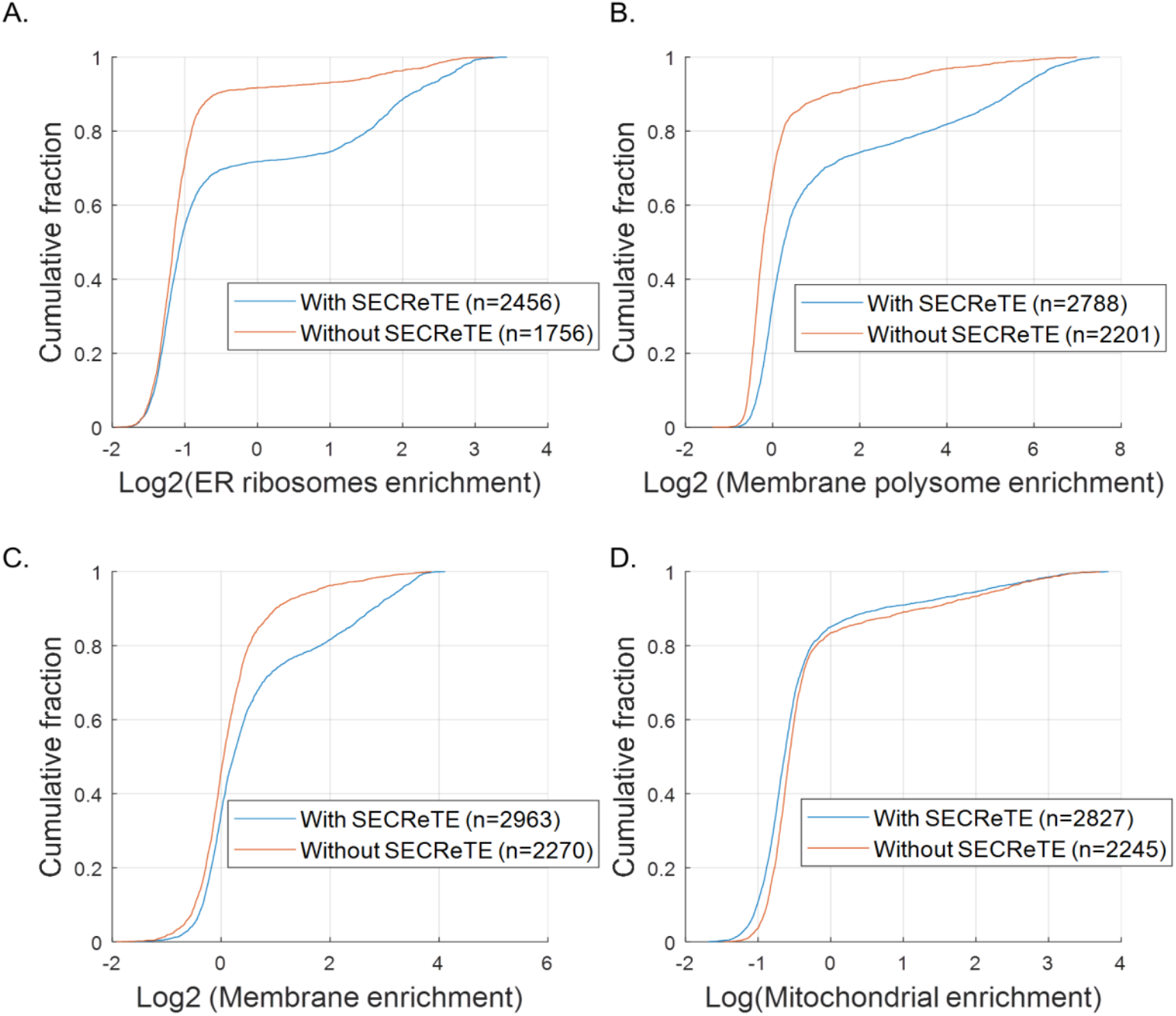
SECReTE-containing transcripts are more enriched in ER, but not mitochondrial, fractions. Cumulative distribution of mRNA enrichment was plotted separately for SECReTE-containing transcripts and transcripts lacking SECReTE. **(A) Transcripts containing SECReTE are more abundant on ER-bound ribosomes.** A plot of the enrichment data obtained from proximity-specific ribosome profiling with BirA-Ubc6 (2min; cycloheximide - CHX)^23^. **(B) Transcripts with SECReTE are more abundant on membranal polysomes.** A plot of the enrichment data obtained from the ribosome profiling of polysomes extracted from the membrane fraction of yeast cells^24^. **(C) Transcripts with SECReTE are more abundant in the membrane fraction.** A plot of the enrichment data obtained from RNA-seq analysis of the membrane fraction of yeast cells^24^. **(D) Transcripts with SECReTE are not abundant on mitochondrial ribosomes.** A plot of the enrichment data obtained from proximity-specific ribosome profiling with BirA-Om45 (2min; CHX)^23^.

**Figure S2.**
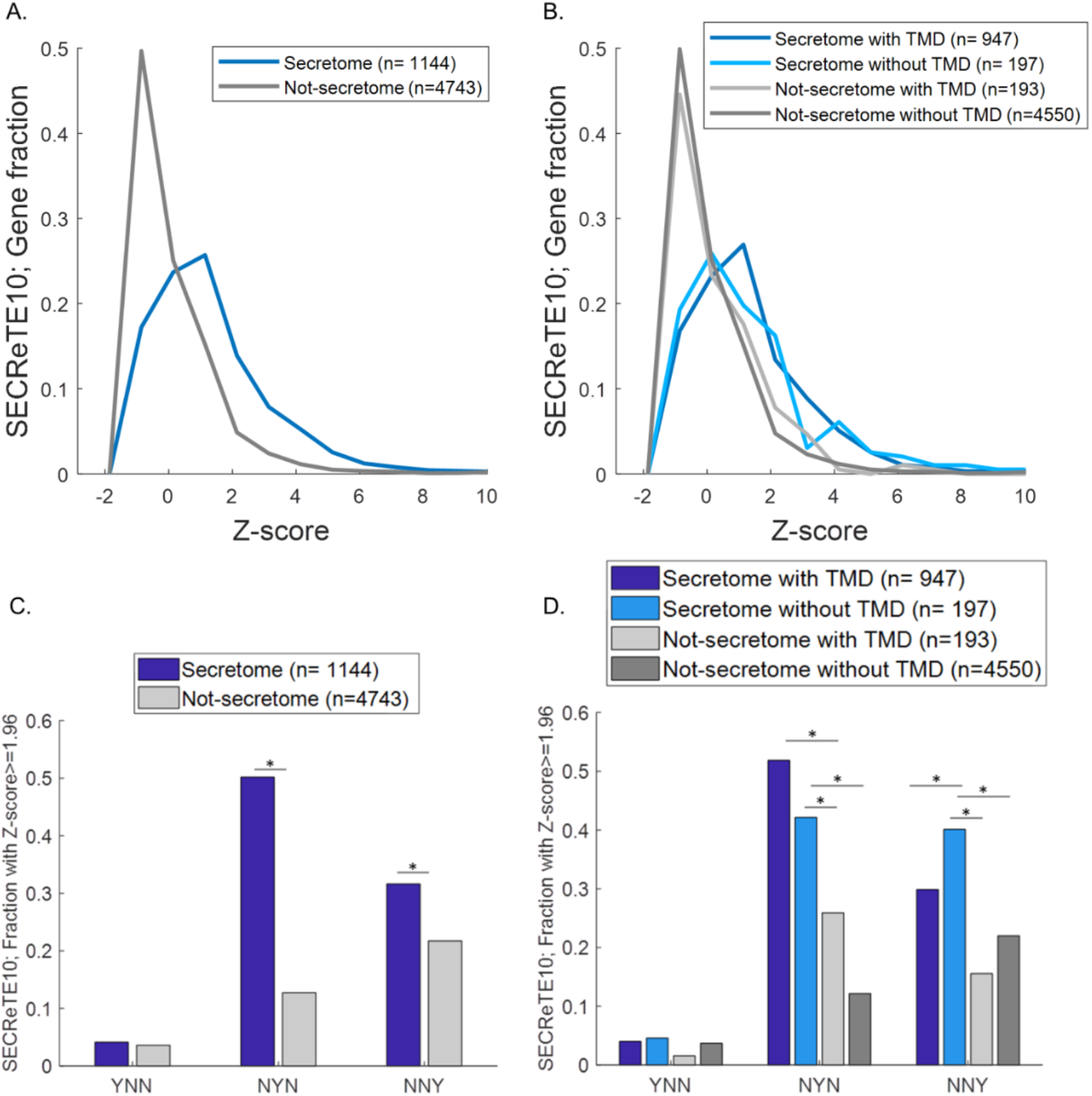
SECReTE abundance is not dependent on codon composition. Permutation analysis was conducted to evaluate the dependency of SECReTE on codon usage. To do that, codon composition was kept and sequences were randomly reshuffled 1000 times. The Z-score was calculated for each gene to assess the probability of the SECReTE10 to appear randomly (for Z-score calculation, see Materials and Methods). The higher the Z-score the less likely it is for SECReTE to appear randomly. **(A) SECReTE enrichment in secretome-encoding mRNAs is independent of codon composition.** Distribution plots of Z-scores show higher values for mRNAs encoding secretome proteins than for non-secretome proteins. **(B) SECReTE enrichment in mRNAs encoding both soluble and membranal secretome transcripts is independent of codon composition.** Distribution plots of Z-scores show higher values for mRNAs encoding secretome proteins (mSMPs; either with or without a TMD) than for non-secretome proteins (*i.e.* with or without a TMD). **(C) SECReTE enrichment in the second and third position of the codon is independent of codon usage.** The fraction of significant Z-scores (*i.e.* ≥1.96) is larger for mRNAs encoding secretome proteins than for non-secretome proteins. **(D) SECReTE enrichment in the second and third position of the codon is independent of both codon usage and TMD presence.** The fraction of significant Z-scores (*i.e.* ≥1.96) is larger for mRNAs encoding secretome proteins than for non-secretome proteins, either with or without a TMD.

**Figure S3.**
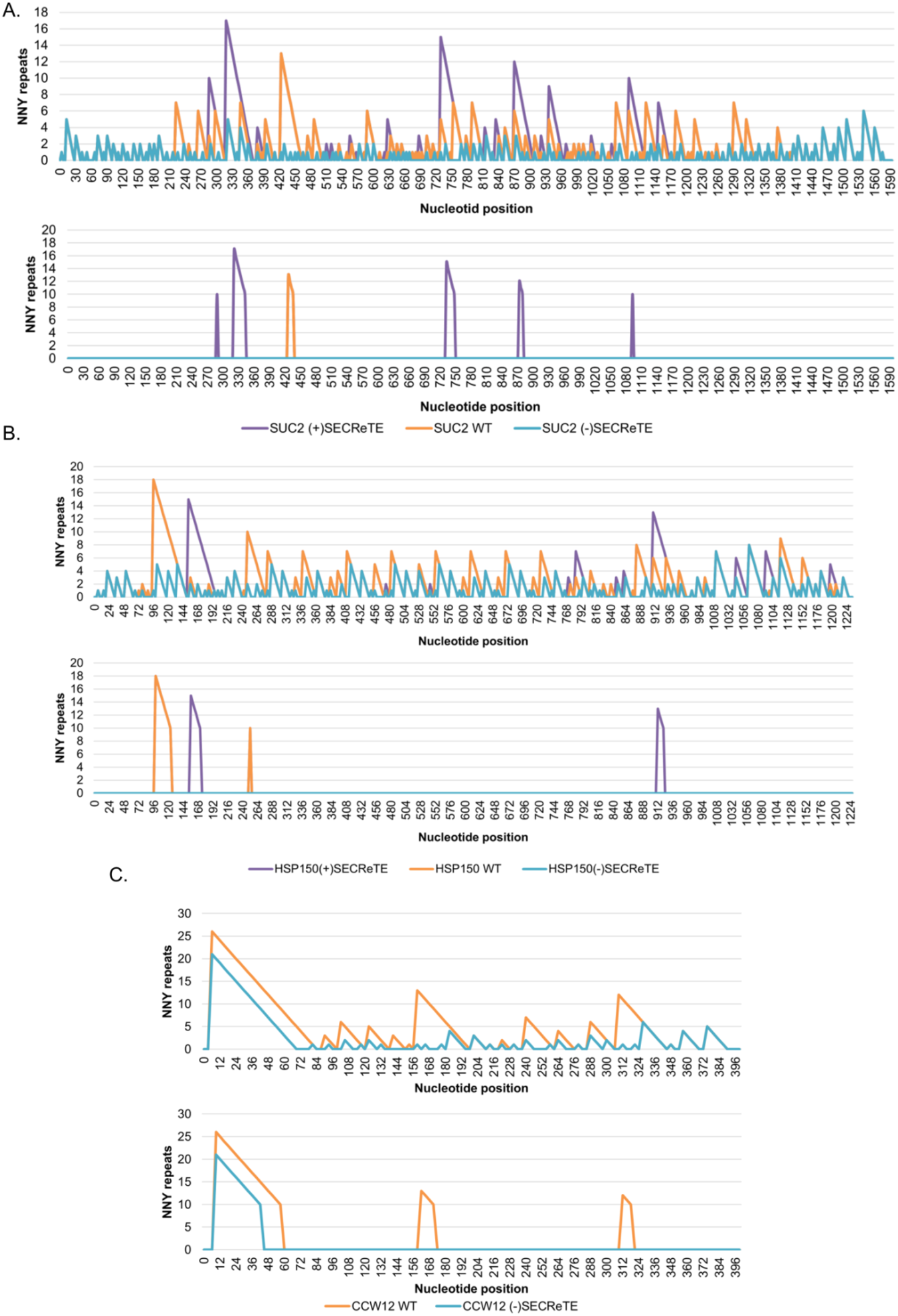
Illustration of SECReTE and SECReTE mutations in *SUC2, HSP150*, and *CCW12*. Graphs compare the number of *NNY* repeats found along the length of the gene either with (lower schematics) or without using a threshold of 10 consecutive *NNY* repeats (upper schematics) in the native and mutant SECReTE genes. **(A)** *SUC2*. **(B)** *HSP150*. **(C)** *CCW12*.

**Figure S4.**
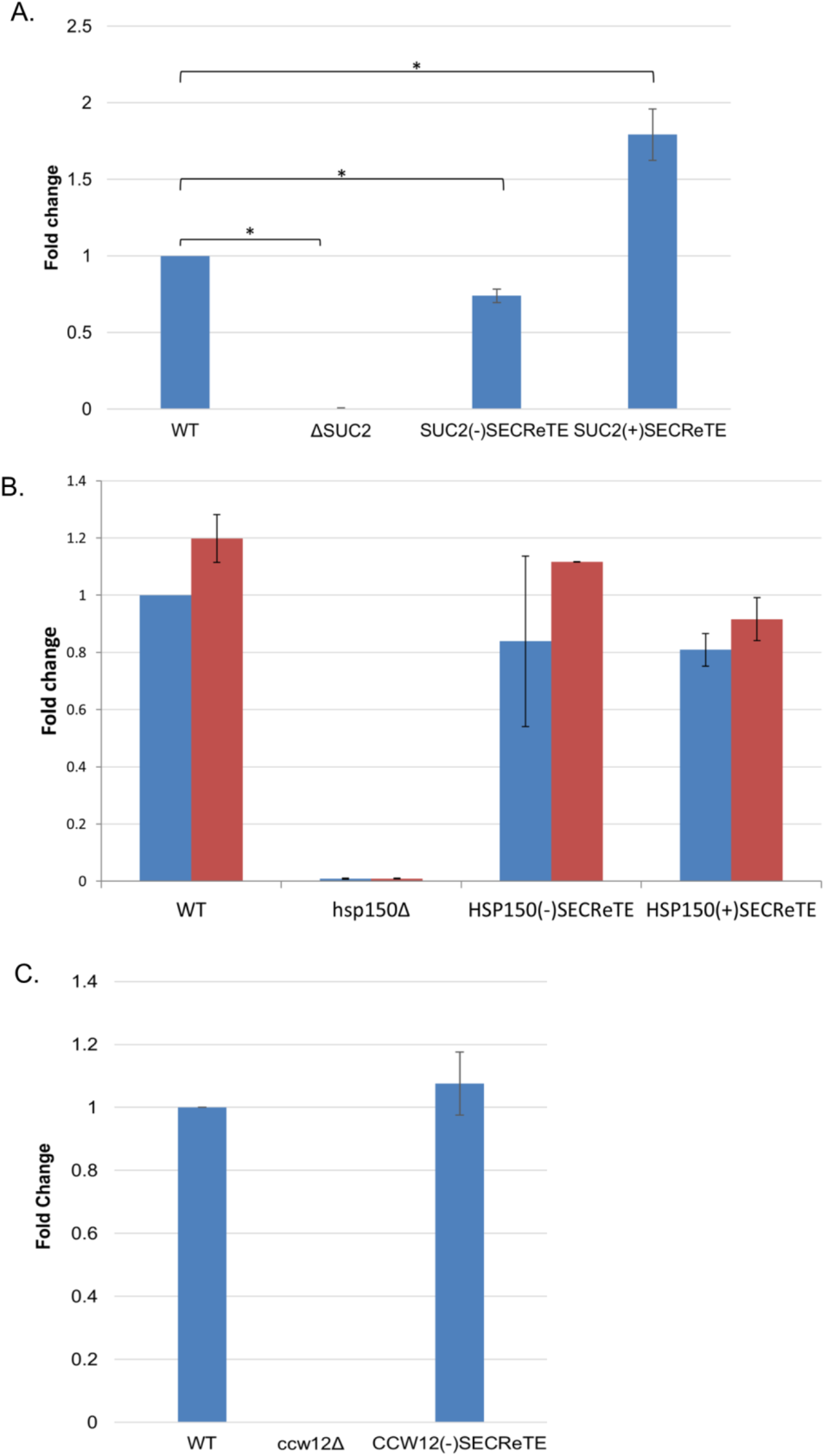
Mutations in SECReTE do not necessarily affect mRNA levels. mRNA levels of native or mutant *SUC2, CCW12*, and *HSP150* in the indicated strains were quantified by qRT-PCR. Fold-change was calculated relative to WT levels. **(A) *SUC2* mRNA levels are altered by SECReTE mutation.** Cells were grown to mid-log phase on SC medium containing 2% glucose at 30°C prior to shifting cells to low glucose medium for 1.5hrs. After harvesting and RNA extraction, primers used for amplifying the long transcript of *SUC2*, which encodes the secreted protein. Primers for actin were used for normalization. *SUC2*(-)SECReTE cells exhibited lower *SUC2* mRNA levels than WT, while *SUC2*(+)SECReTE cells yielded higher levels. Error bars represent the standard deviation of three biological repeats. **(B) *CCW12* mRNA levels are not altered by SECReTE mutation.** Cells were grown to mid-log phase onYPD medium at 30°C prior to harvesting and RNA extraction. Primers used for amplifying *UBC6* were used for normalization. *CCW12* mRNA levels were not significantly changed as a result of SECReTE alterations. **(C) *HSP150* mRNA levels are not altered by SECReTE mutation.** Yeast strains were grown to mid-log phase at either 26°C or 37°C on YPD medium prior to harvesting and RNA extraction. *UBC6* was used for normalization. *HSP150* mRNA levels were not significantly changed as a result of SECReTE alterations.

**Figure S5.**
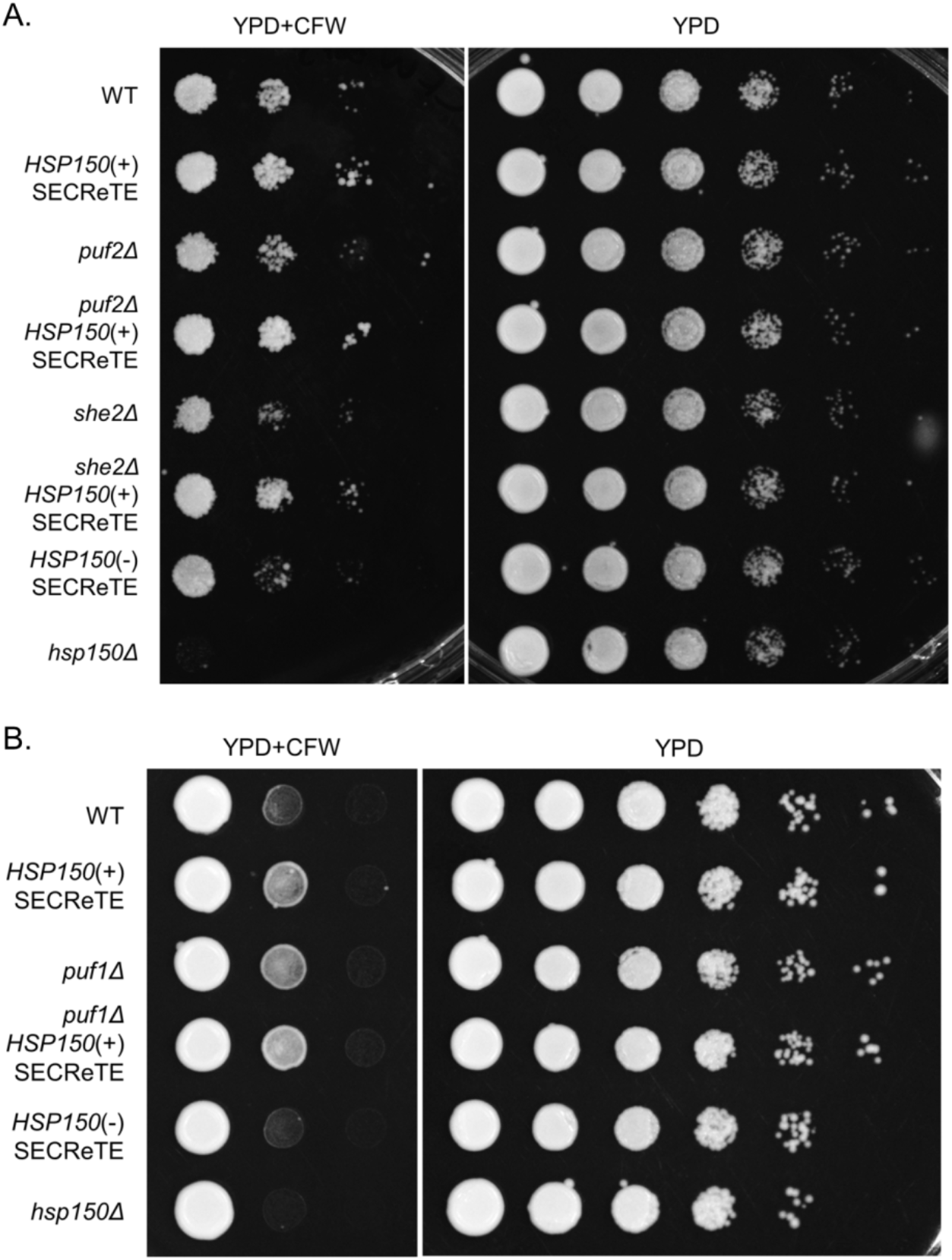
Identification of potential SECReTE-binding proteins. WT cells and either WT or *HSP150*(+)SECReTE cells deleted for genes encoding the indicated RBPs [*e.g.* Puf2, She2 (A) and Puf1(B)] were grown to mid-log phase on YPD at 30°C, prior to serial dilution and plating onto either solid YPD medium or YPD containing CFW. Yeast were grown 2 days prior to photo-documentation.

**Table S1.** Yeast strains used in this study.

| **Name** | **Genotype** | **Source** |
| --- | --- | --- |
| WT BY4741 | *MATa his3Δ1 leu2Δ0 met15Δ0 ura3Δ0* | Euroscarf |
| *suc2Δ* | *MATa his3Δ1 leu2Δ0 met15Δ0 ura3Δ0 suc2Δ::natMX* | This study |
| *SUC2(-)SECRete* | *MATa his3Δ1 leu2Δ0 met15Δ0 ura3Δ0 SUC2::SUC2(-)SECRete* | This study |
| *SUC2(+)*SECRete | *MATa his3Δ1 leu2Δ0 met15Δ0 ura3Δ0 SUC2::SUC2(+)*SECRete | This study |
| *hsp150Δ* | *MATa his3Δ1 leu2Δ0 met15Δ0 ura3Δ0 hsp150Δ::natMX* | This study |
| *hsp150(-)*SECRete | *MATa his3Δ1 leu2Δ0 met15Δ0 ura3Δ0 HSP150::HSP150(-)*SECRete | This study |
| *hsp150(+)*SECRete | *MATa his3Δ1 leu2Δ0 met15Δ0 ura3Δ0 HSP150::HSP150(+)*SECRete | This study |
| *ccw12Δ* | *MATa his3Δ1 leu2Δ0 met15Δ0 ura3Δ0 ccw12Δ::natMX* | This study |
| *CCW12(-)*SECRete | *MATa his3Δ1 leu2Δ0 met15Δ0 ura3Δ0 CCW12::CCW12(-)*SECRete | This study |
| *she2Δ* | *MATa his3Δ1 leu2Δ0 met15Δ0 ura3Δ0 she2Δ::natMX* | This study |
| *she2Δ*<br>*HSP150(+)*SECRete | *MATa his3Δ1 leu2Δ0 met15Δ0 ura3Δ0 she2Δ::natMX*<br>*HSP150::HSP150(+)*SECRete | This study |
| *puf2Δ* | *MATa his3Δ1 leu2Δ0 met15Δ0 ura3Δ0 puf2Δ::natMX* | This study |
| *puf2Δ*<br>*HSP150(+)*SECRete | *MATa his3Δ1 leu2Δ0 met15Δ0 ura3Δ0 puf2Δ::natMX*<br>*HSP150::HSP150(+)*SECRete | This study |
| *puf1Δ* | *MATa his3Δ1 leu2Δ0 met15Δ0 ura3Δ0 puf1Δ::natMX* | This study |
| *puf1Δ*<br>*HSP150(+)*SECRete | *MATa his3Δ1 leu2Δ0 met15Δ0 ura3Δ0 puf1Δ::natMX*<br>*HSP150::HSP150(+)*SECRete | This study |
| *whi3Δ* | *MATa his3Δ1 leu2Δ0 met15Δ0 ura3Δ0 whi3Δ::natMX* | This study |
| *whi3Δ*<br>*HSP150(+)*SECRete | *MATa his3Δ1 leu2Δ0 met15Δ0 ura3Δ0 whi3Δ::natMX*<br>*HSP150::HSP150(+)*SECRete | This study |
| *khd1Δ* | *MATa his3Δ1 leu2Δ0 met15Δ0 ura3Δ0 khd1Δ::natMX* | This study |
| *khd1Δ*<br>*HSP150(+)*SECRete | *MATa his3Δ1 leu2Δ0 met15Δ0 ura3Δ0 khd1Δ::natMX*<br>*HSP150::HSP150(+)*SECRete | This study |

**Table S2.**
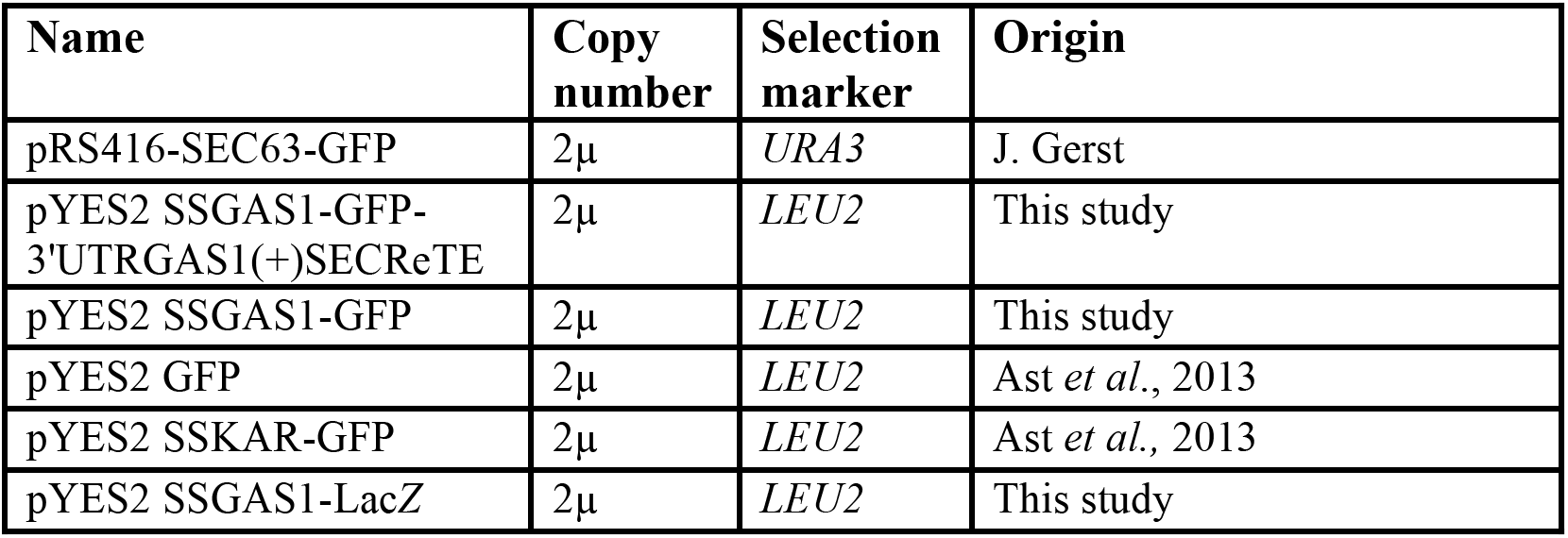
Plasmids used in this study.

| Name | Copy number | Selection marker | Origin |
| --- | --- | --- | --- |
| pRS416-SEC63-GFP | 2μ | *URA3* | J. Gerst |
| pYES2 SSGAS1-GFP-3'UTRGAS1(+)*SECR*eTE | 2μ | *LEU2* | This study |
| pYES2 SSGAS1-GFP | 2μ | *LEU2* | This study |
| pYES2 GFP | 2μ | *LEU2* | Ast *et al.*, 2013 |
| pYES2 SSKAR-GFP | 2μ | *LEU2* | Ast *et al.*, 2013 |
| pYES2 SSGAS1-LacZ | 2μ | *LEU2* | This study |

**Table S3.** SECReTE mutation.

| Name | Length | SECR*eTE* number and codon position |  |  | Free energy | CAI |
| --- | --- | --- | --- | --- | --- | --- |
|  |  | *YNN* | *NYN* | *NNY* |  |  |
| *SUC2* | 1599 | 0 | 0 | 1 | -414.8 | 0.79 |
| *SUC2(+)**SECR*eTE | 1599 | 0 | 0 | 6 | -405.7 | 0.8 |
| *SUC2(-)**SECR*eTE | 1599 | 0 | 0 | 0 | -408.7 | 0.72 |
| *HSP150<sub>WT</sub>* | 1242 | 0 | 4 | 2 | -312.7 | 0.83 |
| *HSP150(+)**SECR*eTE | 1242 | 0 | 4 | 4 | -297 | 0.84 |
| *HSP150(-)**SECR*eTE | 1242 | 0 | 4 | 0 | -358.5 | 0.8 |
| *CCW12<sub>WT</sub>* | 402 | 0 | 3 | 3 | -85.3 | 0.83 |
| *CCW12(-)**SECR*eTE | 402 | 0 | 3 | 1 | -106.9 | 0.8 |

## References

1. Martin K. C. & Ephrussi A. mRNA Localization: Gene Expression in the Spatial Dimension. Cell 136, 719-730 (2009).

2. Buxbaum, A. R., Haimovich, G. & Singer, R. H. In the right place at the right time: visualizing and understanding mRNA localization. Nat. Rev. Mol. Cell Biol. 16, 95-109 (2015).

3. Blobel G. & Dobberstein B. Transfer of proteins across membranes. I. Presence of proteolytically processed and unprocessed nascent immunoglobulin light chains on membrane bound ribosomes of murine myeloma. J. Cell Biol. 67, 835-851 (1975).

4. Gilmore, R., Blobel, G. & Walter, P. Protein translocation across the endoplasmic reticulum.I. Detection in the Microsomal Membrane of a Receptor for the Signal Recognition Particle. J. Cell Biol. 95, 463-469 (1982).

5. Walter P. & Blobel G. Translocation of proteins across membranes III. Signal recognition protein (SRP) causes signal sequence-dependent and site specific arrest of chain elongation that is released by microsomal membranes. J. Cell Biol. 91, 557-561 (1981).

6. Schwartz T. U. Origins and evolution of cotranslational transport to the ER. Adv. Exp. Med. Biol. 607, 52-60 (2007).

7. Saraogi I. & Shan S. Molecular mechanism of co-translational protein targeting by the signal recognition particle. Traffic 12, 535-542 (2011).

8. Kraut-Cohen J. & Gerst J. E. Addressing mRNAs to the ER: cis sequences act up! Trends Biochem. Sci. 35, 459-469 (2010).

9. Weis, B. L., Schleiff, E. & Zerges, W. Protein targeting to subcellular organelles via mRNA localization. Biochim. Biophys. Acta - Mol. Cell Res. 1833, 260-273 (2013).

10. Mutka S. C. & Walter P. Multifaceted Physiological Response Allows Yeast to Adapt to the Loss of the Signal Recognition Particle-dependent Protein-targeting Pathway. Mol. Biol. Cell 12, 577-588 (2001).

11. Ren Y.-G. et al. Differential regulation of the TRAIL death receptors DR4 and DR5 by the signal recognition particle. Mol. Biol. Cell 15, 5064-5074 (2004).

12. Diehn, M., Eisen, M. B., Botstein, D. & Brown, P. O. Large-scale identification of secreted and membrane-associated gene products using DNA microarrays. Nat. Genet. 25, 58-62 (2000).

13. Lerner R. S. et al. Partitioning and translation of mRNAs encoding soluble proteins on membrane-bound ribosomes. RNA 9, 1123-1137 (2003).

14. Pyhtila B. et al. Signal sequence - and translation-independent mRNA localization to the endoplasmic reticulum. RNA 14, 445-453 (2008).

15. Reid D. W. & Nicchitta C. V. Primary role for endoplasmic reticulum-bound ribosomes in cellular translation identified by ribosome profiling. J. Biol. Chem. 287, 5518-5527 (2012).

16. Jagannathan S. et al. De novo translation initiation on membrane-bound ribosomes as a mechanism for localization of cytosolic protein mRNAs to the endoplasmic reticulum. RNA 20, 1489-1498 (2014).

17. Chen, Q., Jagannathan, S., Reid, D. W., Zheng, T. & Nicchitta, C. V. Hierarchical regulation of mRNA partitioning between the cytoplasm and the endoplasmic reticulum of mammalian cells. Mol. Biol. Cell 22, 2646-2658 (2011).

18. Kraut-Cohen J. et al. Translation - and SRP-independent mRNA targeting to the endoplasmic reticulum in the yeast Saccharomyces cerevisiae. Mol. Biol. Cell 24, 3069-84 (2013).

19. Ast, T., Cohen, G. & Schuldiner, M. A network of cytosolic factors targets SRP-independent proteins to the endoplasmic reticulum. Cell 152, 1134-1145 (2013).

20. Aviram N. et al. The SND proteins constitute an alternative targeting route to the endoplasmic reticulum. Nature 540, 134-138 (2016).

21. Johnson, N., Powis, K. & High, S. Post-translational translocation into the endoplasmic reticulum. Biochim. Biophys. Acta - Mol. Cell Res. 1833, 2403-2409 (2013).

22. Jan, C. H., Williams, C. C. & Weissman, J. S. Principles of ER cotranslational translocation revealed by proximity-specific ribosome profiling. Science 346, 748-751 (2014).

23. Chartron, J. W., Hunt, K. C. L. & Frydman, J. Cotranslational signal-independent SRP preloading during membrane targeting. Nature 536, 224-228 (2016).

24. Shahbabian K. & Chartrand P. Control of cytoplasmic mRNA localization. Cell. Mol. Life Sci. 69, 535-552 (2012).

25. Hamilton R. S. & Davis I. Identifying and searching for conserved RNA localisation signals. Methods Mol. Biol. 714, 447-466 (2011).

26. Polyansky A.a, Hlevnjak, M. & Zagrovic B. Analogue encoding of physicochemical properties of proteins in their cognate messenger RNAs. Nat. Commun. 4, 2784 (2013).

27. Cui X. a. & Palazzo A. F. Localization of mRNAs to the endoplasmic reticulum. Wiley Interdiscip. Rev. RNA 5, 481-492 (2014).

28. Palazzo A. F. et al. The signal sequence coding region promotes nuclear export of mRNA. PLoS Biol. 5, 2862-2874 (2007).

29. Wolfenden R. V, Cullis, P. M. & Southgate C. C. Water, protein folding, and the genetic code. Science 206, 575-577 (1979).

30. Prilusky J. & Bibi E. Studying membrane proteins through the eyes of the genetic code revealed a strong uracil bias in their coding mRNAs. Proc. Natl. Acad. Sci. U. S. A. 106, 6662–6666 (2009).

31. Haim, L., Zipor, G., Aronov, S. & Gerst, J. E. A genomic integration method to visualize localization of endogenous mRNAs in living yeast. Nat. Methods 4, 409-412 (2007).

32. Storici F. & Resnick M. A. The Delitto Perfetto Approach to In Vivo Site-Directed Mutagenesis and Chromosome Rearrangements with Synthetic Oligonucleotides in Yeast. Methods Enzymol. 409, 329-345 (2006).

33. Ye J. et al. Primer-BLAST: A tool to design target-specific primers for polymerase chain reaction. BMC Bioinformatics 13, 134 (2012).

34. Ram A. F. J. & Klis F. M. Identification of fungal cell wall mutants using susceptibility assays based on Calcofluor white and Congo red. Nat. Protoc. 1, 2253-2256 (2006).

35. Zhang T. et al. An improved method for whole protein extraction from yeast Saccharomyces cerevisiae. Yeast 28, 795-798 (2011).

36. Goldstein A. & Lampen J. O. Beta-D-fructofuranoside fructohydrolase from yeast. Methods Enzymol. 42, 504-511 (1975).

37. Novick P. & Schekman R. Secretion and cell-surface growth are blocked in a temperature-sensitive mutant of Saccharomyces cerevisiae. Proc. Natl. Acad. Sci. 76, 1858-1862 (1979).

38. Troy AA, & Harkness, H. T. A. Simplified Method for Measuring Secreted Invertase Activity in Saccharomyces cerevisiae. Biochem. Pharmacol. Open Access 3, 151 (2014).

39. Bailey T. L. et al. MEME SUITE: tools for motif discovery and searching. Nucleic Acids Res. 37, W202-W208 (2009).

40. Sharp P. M. & Li, W. H. The codon Adaptation Index-a measure of directional synonymous codon usage bias, and its potential applications. Nucleic Acids Res. 15, 1281-95 (1987).

41. Denic V. A portrait of the GET pathway as a surprisingly complicated young man. Trends Biochem. Sci. 37, 411-417 (2012).

42. Stefanovic S. & Hegde R. S. Identification of a Targeting Factor for Posttranslational Membrane Protein Insertion into the ER. Cell 128, 1147-1159 (2007).

43. Hsu, P.-H., Chiang, P.-C., Liu, C.-H. & Chang, Y.-W. Characterization of Cell Wall Proteins in Saccharomyces cerevisiae Clinical Isolates Elucidates Hsp150p in Virulence. PLoS One 10, e0135174 (2015).

44. Russo, P., Simonen, M., Uimari, A., Teesalu, T. & Makarow, M. Dual regulation by heat and nutrient stress of the yeast HSP150 gene encoding a secretory glycoprotein. Mol. Gen. Genet. 239, 273-80 (1993).

45. Russo, P., Kalkkinen, N., Sareneva, H., Paakkola, J. & Makarow, M. A heat shock gene from Saccharomyces cerevisiae encoding a secretory glycoprotein. Proc. Natl. Acad. Sci. U. S. A. 89, 3671-5 (1992).

46. Ragni, E., Sipiczki, M. & Strahl, S. Characterization of Ccw12p, a major key player in cell wall stability of Saccharomyces cerevisiae. Yeast 24, 309-319 (2007).

47. Ragni E. et al. The genetic interaction network of CCW12, a Saccharomyces cerevisiae gene required for cell wall integrity during budding and formation of mating projections. BMC Genomics 12, 107 (2011).

48. Colomina N., Ferrezuelo F., Wang H., Aldea M. & Garí, E. Whi3, a developmental regulator of budding yeast, binds a large set of mRNAs functionally related to the endoplasmic reticulum. J. Biol. Chem. 283, 28670-28679 (2008).

49. Hasegawa, Y., Irie, K. & Gerber, A. P. Distinct roles for Khd1p in the localization and expression of bud-localized mRNAs in yeast. RNA 14, 2333-2347 (2008).

50. Hogan, D. J., Riordan, D. P., Gerber, A. P., Herschlag, D. & Brown, P. O. Diverse RNA-binding proteins interact with functionally related sets of RNAs, suggesting an extensive regulatory system. PLoS Biol. 6, 2297-2313 (2008).

51. Reid D. W. & Nicchitta C. V. Diversity and selectivity in mRNA translation on the endoplasmic reticulum. Nat. Rev. Neurosci. 16, 221-231 (2015).

52. Cui, X. A., Zhang, H., Palazzo, A. F., Fugate, R. D. & Reichlin, M. p180 Promotes the Ribosome-Independent Localization of a Subset of mRNA to the Endoplasmic Reticulum. PLoS Biol. 10, e1001336 (2012).

53. Reid D. W. & Nicchitta C. V. Comment on ‘Principles of ER cotranslational translocation revealed by proximity-specific ribosome profiling’. Science 348, 1217 (2015).

54. Jan, C. H., Williams, C. C. & Weissman, J. S. Response to Comment on ‘Principles of ER cotranslational translocation revealed by proximity-specific ribosome profiling’. Science 348, 1217 (2015).

55. Gerst J. E. Message on the web: mRNA and ER co-trafficking. Trends Cell Biol. 18, 68-76 (2008).

56. Kejiou N. S. & Palazzo A. F. mRNA localization as a rheostat to regulate subcellular gene expression. Wiley Interdiscip. Rev. RNA 8, e1416 (2017).

57. Chin A. & Lécuyer E. RNA localization: Making its way to the center stage. Biochim. Biophys. Acta 1861, 2956-2970 (2017).

58. Saint-Georges Y. et al. Yeast Mitochondrial Biogenesis: A Role for the PUF RNA-Binding Protein Puf3p in mRNA Localization. PLoS One 3, e2293 (2008).

59. Houshmandi S. S. & Olivas W. M. Yeast Puf3 mutants reveal the complexity of Puf-RNA binding and identify a loop required for regulation of mRNA decay. RNA 11, 1655-66 (2005).

60. Olivas W. & Parker R. The Puf3 protein is a transcript-specific regulator of mRNA degradation in yeast. EMBO J. 19, 6602-11 (2000).

61. Cai, Y., Futcher, B., Waern, K., Shou, C. & Raha, D. Effects of the Yeast RNA-Binding Protein Whi3 on the Half-Life and Abundance of CLN3 mRNA and Other Targets. PLoS One 8, e84630 (2013).

62. Vergés, E., Colomina, N., Garí, E., Gallego, C. & Aldea, M. Cyclin Cln3 Is Retained at the ER and Released by the J Chaperone Ydj1 in Late G1 to Trigger Cell Cycle Entry. Mol. Cell 26, 649-662 (2007).

63. Ito, W., Li, X., Irie, K., Mizuno, T. & Irie, K. RNA-Binding Protein Khd1 and Ccr4 Deadenylase Play Overlapping Roles in the Cell Wall Integrity Pathway in Saccharomyces cerevisiae. Eukaryot. Cell 10, 1340-1347 (2011).

64. Irie K. et al. The Khd1 protein, which has three KH RNA-binding motifs, is required for proper localization of ASH1 mRNA in yeast. EMBO J. 21, 1158-67 (2002).

65. Aronov S. et al. mRNAs Encoding Polarity and Exocytosis Factors Are Cotransported with the Cortical Endoplasmic Reticulum to the Incipient Bud in Saccharomyces cerevisiae. Mol. Cell. Biol. 27, 3441-3455 (2007).

66. Schmid, M., Jaedicke, A., Du, T.-G. & Jansen, R.-P. Coordination of Endoplasmic Reticulum and mRNA Localization to the Yeast Bud. Curr. Biol. 16, 1538-1543 (2006).

67. Tang H. et al. Engineering vesicle trafficking improves the extracellular activity and surface display efficiency of cellulases in Saccharomyces cerevisiae. Biotechnol. Biofuels 10, 53 (2017).

68. Roohvand, F., Shokri, M., Abdollahpour-Alitappeh, M. & Ehsani, P. Biomedical applications of yeast-a patent view, part one: yeasts as workhorses for the production of therapeutics and vaccines. Expert Opin. Ther. Pat. 27, 929-951 (2017).

